# The Dorsomedial Prefrontal Cortex Uses Reward Predictions to Regulate How Rewards Are Pursued

**DOI:** 10.64898/2026.08.29.748037

**Authors:** Briac Halbout, Collin Hutson, Neha Ramiah, Gabriela Arias, Nitya Naik, Kate M. Wassum, Sean B. Ostlund

**Affiliations:** Department of Psychology, Cal State Long Beach, Long Beach, CA 90824; Department of Anesthesiology and Perioperative Care, UCI, Irvine, CA, 92697; Department of Psychology and Brain Research Institute, UCLA, Los Angeles, California 90095; Department of Neurobiology and Behavior and Irvine Center for Addiction Neuroscience, UCI, Irvine, CA, 92697

**Keywords:** prefrontal cortex, Pavlovian-instrumental transfer, reward expectancy, cognitive control, fiber photometry, chemogenetics

## Abstract

Reward-predictive cues regulate not only whether but how rewards are pursued, and their influence depends on the strength of their predictive relationship with reward. Whereas weak cues signaling sparse or uncertain reward invigorate instrumental reward seeking, cues signaling imminent reward produce little net invigoration of instrumental behavior and instead promote conditioned goal-approach. We propose that this reflects two opposing influences: an automatic motivational impulse to seek reward, and a top-down, expectancy-dependent control process that constrains it. On this view, the muted or even suppressive effect of imminent-reward cues on instrumental reward seeking reflects an active constraint on motivation rather than its absence. Consistent with this account, we found that selectively devaluing the predicted reward unmasked a latent motivational influence. Cues for imminent reward now invigorated rather than constrained instrumental seeking, suggesting the involvement of a goal-directed regulatory process. Separately, shifting rats from hunger to general satiety abolished the cue-specific regulation of instrumental seeking. We then tested whether the dorsomedial prefrontal cortex (dmPFC) mediates this control. Bulk calcium recordings revealed phasic dmPFC activity that encoded predicted reward probability, dipped when reward was omitted, and covaried with instrumental performance in a cue-dependent manner. Moreover, chemogenetic dmPFC inhibition disrupted the cue-specific regulation of instrumental seeking, leaving rats unable to use these predictions to determine how vigorously to press. These findings identify the dmPFC as a substrate for top-down, expectancy-dependent control over reward pursuit and suggest that its dysfunction may contribute to impulsive or maladaptive reward seeking.

**Significance Statement:** Cues for imminent reward availability normally suppress the impulse to perform instrumental reward-seeking actions, redirecting behavior toward goal-approach responses that facilitate reward retrieval. This flexible regulatory process depends on a real-time appraisal of predicted reward value and is lifted when that reward is devalued, allowing such cues to invigorate instrumental behavior. Here we identify the dorsomedial prefrontal cortex (dmPFC) as a critical substrate for this process. Using fiber photometry and chemogenetic inhibition, we show that dmPFC activity tracks these cue-elicited reward predictions and is required for their regulatory influence over instrumental reward seeking, but not over conditioned goal-approach. These findings illuminate a prefrontal mechanism for top-down motivational control with potential relevance to impulsive reward seeking.

## Introduction

Reward-predictive cues shape not only whether animals pursue reward but how they do so (Konorski, 1967; Bindra, 1974; Timberlake, 1994). Their influence over instrumental reward seeking depends on the strength and nature of their predictive relationship with reward (Ostlund and Marshall, 2021). For instance, weak cues that signal sparse or distal reward availability tend to invigorate instrumental reward seeking (Estes, 1943; Corbit and Balleine, 2015). In contrast, strong cues that signal a high probability of imminent reward delivery lack this excitatory effect and may even suppress instrumental performance (Azrin and Hake, 1969; Miczek and Grossman, 1971; Crombag et al., 2008; Marshall et al., 2020; Ostlund and Marshall, 2021). Such cues instead preferentially evoke more intensive conditioned goal-approach (checking the anticipated site of reward delivery). This ability to regulate reward pursuit as reward expectancy changes is a fundamental characteristic of motivation control, and when lost may contribute to impulsive reward seeking (Timberlake, 1988; Robinson et al., 2018; Ostlund and Marshall, 2021). Although much is known about the neural systems mediating the excitatory motivational influence of cues on instrumental reward seeking (Corbit and Balleine, 2015), the mechanisms that constrain instrumental performance when imminent reward is predicted remain to be elucidated.

The current study investigated the role of the dorsomedial prefrontal cortex (dmPFC; comprising interconnected anterior cingulate and prelimbic subregions) in this phenomenon. The dmPFC is known to play a critical role in exerting flexible control over reward pursuit and resolving response conflict (Shenhav et al., 2013; Sharpe et al., 2019; McLaughlin et al., 2021; Rudebeck and Izquierdo, 2022; Clairis and Lopez-Persem, 2023). However, its role in cue-motivated behavior remains unclear (Howland et al., 2022). Several studies have investigated dmPFC involvement in the Pavlovian-to-instrumental transfer (PIT) task, which is conventionally used to assay the tendency for a weakly predictive cue to motivate an independently trained instrumental action (Estes, 1943; Corbit and Balleine, 2015). Such studies have consistently found that this capacity is spared following lesions or inhibition of the dmPFC (Cardinal et al., 2003; Corbit and Balleine, 2003; Halbout et al., 2022). However, chemogenetic dmPFC stimulation suppresses the tendency for weak cues to motivate instrumental reward seeking while leaving conditioned goal-approach intact (Halbout et al., 2022). Moreover, dmPFC neurons have been shown to phasically encode cue-triggered instrumental actions and conditioned goal approach in overlapping populations, and do so with a predominantly inhibitory signal that covaries with PIT task performance (Homayoun and Moghaddam, 2009). These latter findings suggest the dmPFC may be involved in regulating the expression of cue-motivated reward seeking.

The current study tests the hypothesis that the dmPFC uses cue-elicited reward predictions to shape the way rewards are pursued, supporting flexible control over instrumental reward seeking when it is more adaptive to approach the goal location. We examined this possibility using a variant of the PIT task in which cues signal a high (100%) or low (30%) probability of imminent reward delivery (10-s cue-reward interval), contrasting their effects on instrumental reward seeking and conditioned goal approach (Marshall et al., 2020; Ostlund and Marshall, 2021; Malvaez et al., 2026). We first tested a central assumption of this account, that cues for imminent reward acquire motivational properties which are actively suppressed when the predicted reward is highly valued, compelling rapid retrieval. We examined if this latent motivational influence could be unmasked by devaluing the predicted reward through conditioned taste aversion learning. We then assessed the impact of a general shift from hunger to satiety, which reduces both the value of the predicted reward and the presumed (latent) excitatory motivational influence of the eliciting cue. Next, we used fiber photometry to record dmPFC Ca^2+^ activity during the task to determine whether it encodes predicted reward probability and its regulatory influence on instrumental reward seeking. Finally, we tested whether chemogenetic dmPFC inhibition disrupts the capacity to use predictive cues to selectively regulate instrumental reward seeking and not conditioned goal-approach.

## Materials and Methods

### Animals

Adult (>75 days) Long-Evans rats (Envigo) were used in this study. Rats were housed in pairs or singly (after surgery) in standard Plexiglas cages on a 12h/12h light/dark cycle and were maintained at ∼85% of their free-feeding bodyweight during behavioral procedures, unless otherwise noted. All experimental procedures involving live animals were approved by the UC Irvine Institutional Animal Care and Use Committee and were in accordance with the National Research Council Guide for the Care and Use of Laboratory Animals.

### Apparatus

Behavioral procedures took place in sound- and light-attenuated Med Associates rat operant chambers. Food pellets (grain-based or banana-flavored purified, 45 mg; BioServ) or sweetened condensed milk solution (SCM; 1 ml of 50% v/v in water) were delivered into a food cup positioned in a recessed port, which was adjacent to the lever. A photobeam detector was used to record food-port entries. Chambers were illuminated during all sessions and a ventilation fan provided continuous background noise (∼60 dB). For the fiber photometry experiment, asymptotic training and the test session were conducted in a similar operant chamber equipped with identical stimulus and response devices but housed in an adjacent room.

### Surgery

Surgeries were performed as in our prior studies (Halbout et al., 2019; Halbout et al., 2022) under aseptic conditions. Rats were anesthetized with isoflurane (2-5% in oxygen at 0.5L/min) and were injected with the nonsteroidal anti-inflammatory carprofen (5 mg/kg; sc) and the antibiotic enrofloxacin (10 mg/kg; sc) before being placed in a stereotaxic frame. After intracranially infusing adeno-associated virus (AAV) vector into the dmPFC and implanting optical or infusion cannulas (see below for details), the incision was closed with sutures or dental acrylic. A second dose of carprofen was administered 24h post-surgery and animals were observed and weighed daily for at least 7d before undergoing behavioral training.

### Drug preparation and administration

Clozapine-n-oxide (CNO) obtained from NIMH (NIMH Chemical Synthesis and Drug Supply Program) was dissolved in 5% DMSO in sterile saline for intraperitoneal administration (1 mL/kg).

### Histology

After behavioral testing, rats received a lethal dose of Euthasol (150 mg/kg sodium pentobarbital, i.p.) before being perfused with PBS followed by 4% paraformaldehyde. Brains were post-fixed with 4% paraformaldehyde, cryoprotected with 30% sucrose, and sectioned into 40-µm coronal sections on a cryostat. To visualize expression, hM4Di, GFP, and GCaMP signals were immunohistochemically amplified using antibodies directed against mCherry (hM4Di) or GFP (GFP, GCaMP). Tissue was first incubated in 3% normal donkey serum in PBS plus Triton X-100 (PBST; 1 h) and then in primary antibodies in PBST at 4°C for 48 h, using rabbit anti-DsRed (mCherry tag; 1:1,000; Clontech; 632496) or mouse anti-GFP (1:1,500; Life Technologies; A- 11120). Sections were then incubated for 2 h at room temperature in fluorescent conjugated secondary antibodies, Alexa Fluor 594 goat anti-rabbit (DsRed; 1:500; Invitrogen; A11037) or Alexa Fluor 488 goat anti-mouse (GFP; 1:500; Invitrogen; A10667). Sections were mounted with mounting medium containing DAPI (Vectashield) and imaged with a 10× objective on a fluorescence microscope (Leica) to validate viral expression.

## Reward devaluation experiment: impact of outcome-specific devaluation on regulation of cue-motivated behavior

### Overview

24 adult male rats were food-deprived and trained on a variant of the PIT task designed to probe the role of outcome-specific reward representations in the regulatory influence of strong reward-predictive cues.

### Food-port training

Rats were given two daily sessions in which 30 food rewards were delivered on a random-time (RT) 60-s schedule into the food port. Banana pellets were delivered in session 1, whereas both SCM and grain pellets were delivered (15 each) in session 2.

### Pavlovian conditioning

Rats then underwent 9 sessions of Pavlovian conditioning, in which two 10-s auditory conditioned stimuli (CS; 3-kHz tone and 10-Hz clicker; 85dB) signaled delivery of distinct reward outcomes (O1 and O2; SCM or grain pellet) at CS offset with either a 100% (n = 12) or 30% (n = 12) probability. In each group, CS-outcome contingencies were counterbalanced. In each session, a 55-s interval preceded the onset of the first CS and trials were separated by a 110-s inter-trial interval (ITI). Each session consisted of 15 trials of each CS alternating in a pseudorandom order (30 trials total per session).

### Instrumental training

Rats then received 9 sessions of instrumental training, during which the lever was continuously inserted into the chamber. In the first session, each lever-press was reinforced (fixed-ratio-1, FR-1) with a banana pellet (O3) with a 1-s post-reward time out period. This session lasted a maximum of 30 min or until 30 pellets had been earned. Additional FR-1 sessions were provided as needed until rats earned all 30 pellets within 30 min. During subsequent sessions, lever pressing was reinforced according to a random-interval (RI) schedule, such that the lever remained available but was inactive for an average of *t* seconds after each reward delivery, with individual *t* values selected from an exponential distribution. The RI schedule was adjusted over days to establish steady and persistent lever-press behavior, with one day of RI-15 (t = 15 s), one day of RI-30 (t = 30 s), and six days of RI-60 (t = 60 s). RI sessions lasted a maximum of 30 minutes or were terminated after rats earned 30 pellets.

### Conditioned taste aversion training

One of the two Pavlovian outcomes (O1 or O2) was then selectively devalued through conditioned taste aversion learning. Individual rats were given 60-min access to O1 or O2 on alternating days in standard housing cages (five sessions with each outcome). During each session, rats received either 20 g of grain pellets in a metal food cup or SCM in a precision drinking bottle. Immediately following each session, rats received an i.p. injection of either LiCl (0.15 M; 20 mL/kg) or sterile saline, so that consumption of one outcome was consistently paired with gastric malaise and consumption of the other with saline (Holland, 2004; Marshall et al., 2023).

### Instrumental retraining and extinction

Rats were given one day of RI-60 instrumental retraining (as above). On the next day, they were given a 30-min instrumental extinction session, in which the lever was continuously available but inactive.

### PIT test

On the next day, rats received a 40-min PIT test session, during which the lever was continuously available but inactive. The test included 10 noncontingent presentations of each of the two 10-s CS in a pseudorandom order (20 trials total). The ITI was 90 s, and a 5-min interval preceded the onset of the first CS (i.e., 3.5 min plus one ITI). No rewards were delivered at test.

## Satiety-shift experiment: impact of a shift from hunger to general satiety on regulation of cue-motivated behavior

### Overview

Adult male (n = 24) and female (n = 16) rats were food-deprived and trained on a probabilistic PIT task, allowing us to assess how a general shift from hunger to satiety affects their ability to flexibly regulate their instrumental performance when presented with weak versus strong reward-paired cues.

### Food-port training

Rats were given two daily 30-min sessions with grain pellets delivered on a RT 60-s schedule.

### Pavlovian conditioning

Rats received 9 sessions of Pavlovian conditioning in which two 10-s CSs (tone and clicker) differentially signaled either a 30% or 100% probability of grain pellet reward delivery at CS offset (CS_30_ and CS_100_, respectively). CS identity was counterbalanced with reward probability (and other experimental variables). Trials began after an initial 55-s interval and were separated by a 110-s ITI and ended after 40 trials (20 with each CS, pseudorandom order).

### Instrumental training

Rats then received 9 sessions of instrumental training, as in the reward devaluation experiment, in which they learned to lever press for grain pellets with one day each of FR-1, RI-15 and RI-30, and six days of RI-60. Sessions ended after 30 minutes or until 30 pellets were earned.

### Pavlovian retraining and extinction

Rats received a session of Pavlovian retraining (as above) and a 30-min instrumental extinction session (lever was continuously available but inactive) on the following day.

### PIT testing

*On the following day,* rats received a PIT test using the same procedure as the reward devaluation experiment but were tested either in state of hunger or general satiety, which was induced by providing them with free access to lab chow in their home cage for 18h prior to testing (i.e., initiated after instrumental extinction session). Food restriction was reinitiated for all rats after the PIT test. After 72h, rats were given three days of Pavlovian retraining and two days of instrumental retraining before undergoing a session of instrumental extinction and a second PIT test in the alternate need state (i.e., rats previously tested hungry were now tested sated, and vice versa), providing a within-subject manipulation of this variable.

## Photometry experiment: bulk calcium recordings from dmPFC neurons during regulation of cue-motivated behavior

### Overview

Adult male rats were used in a fiber photometry experiment to record bulk calcium signaling in dmPFC neurons, in order to characterize neural activity evoked by strong versus weak reward-predictive cues during the probabilistic PIT task. Our primary recording experiment used the same 10-s delay Pavlovian conditioning procedure used in other experiments in the study (n = 9). To explore the influence of reward timing on behavior and dmPFC calcium signaling, we also ran a small cohort (n = 5) trained with a 5-s delay conditioning procedure, in which reward was delivered midway through the 10-s CS presentation. Reward probabilities (100% and 30%) were identical in both groups.

### Surgery

Adult male rats underwent surgery to inject 0.3 µL (0.1 µL/min) of the calcium indicator GCaMP6s (AAV9-CaMKII-GCaMP6s; Addgene 107790-AAV9) into the dmPFC (AP: +2.2, ML: +0.6; DV: −2.8; mm relative to bregma) with a 400-µm diameter optical cannula (Doric Lenses) implanted 0.5 mm above the injection site and secured to the skull with four stainless screws.

### Food-port training

Identical to the satiety-shift experiment.

### Pavlovian conditioning

Identical to the satiety-shift experiment, except that rats received some training (including their last session) in the fiber photometry test chamber to facilitate generalization, and except that rats in the 5-s delay group received reward at a fixed time 5 s after cue onset rather than at cue offset.

### Instrumental training

Identical to the satiety-shift experiment, except the final training session took place in the fiber photometry recording chamber.

### Pavlovian retraining and instrumental extinction

Identical to the satiety-shift experiment, except the final training and extinction sessions took place in the fiber photometry recording chamber to facilitate behavioral generalization.

### PIT testing

Identical to the satiety-shift experiment, except the test took place in the fiber photometry recording chamber.

### Fiber photometry

Recordings were made using a Tucker Davis Technologies (TDT) RZ10x processor and TDT synapse software. LEDs for 465nm and 405nm (isosbestic) excitation light were delivered (modulated at 210 Hz and 330 Hz, respectively) through a low autofluorescence optical patch cable (fiber core diameter 400 µm; Doric Lenses) connected to the implanted optical cannula. Excitation light power was adjusted to between 10-30uW (measured at the fiber tip). Signals were separated by a Doric Lenses fluorescence minicube and were offset by 5mA and demodulated using a 4 Hz lowpass filter. TTL signals encoding behavioral events were transmitted to the rig and aligned to photometry data, which were analyzed offline in a custom Python-based pipeline (see below).

## Chemogenetic inhibition experiment: impact of dmPFC inhibition on regulation of cue-motivated behavior

### Overview

Adult male rats were used to determine whether dmPFC function is necessary for regulating expression of cue-motivated behavior on the probabilistic PIT task. Surgery: Rats received bilateral injections of AAV to express the inhibitory designer receptor hM4Di (AAV5-CaMKIIa-hM4D(Gi)-mCherry; Addgene 50477; n = 14) or GFP alone (AAV5-CaMKIIa-EGFP; Addgene 50469; n = 12) in 6 bilateral sites along the anteroposterior dimension of the dmPFC (+3.2 AP, +/− 0.6 ML, −2.8 DV; +2.2 AP, +/− 0.6 ML, −2.2 DV; +1.2 AP, +/− 0.6 ML, −3.2 DV). Each injection site received 0.3 µL of virus, infused at a rate of 0.1 µL/min.

### Behavioral training

Food-port training, Pavlovian conditioning, and instrumental training, Pavlovian retraining and instrumental extinction procedures were identical to the satiety-shift experiment.

### PIT testing

Rats were pre-treated with 5 mg/kg CNO or vehicle (1 ml/kg of 5% DMSO in sterile saline, i.p.) 30 min prior to the PIT test. This dose has been used previously to chemogenetically inhibit hM4Di-expressing neurons in rats undergoing PIT testing with little or no evidence of nonspecific behavioral effects in hM4Di-free controls (Halbout et al., 2019; Halbout et al., 2022). Each rat was tested four times, twice in each drug condition, with order of treatment counterbalanced across all experimental conditions. Rats received 2 days of instrumental retraining (RI-60s), one day of Pavlovian retraining, and one day of instrumental extinction before each round of retesting. Data were averaged within each drug condition prior to statistical analysis.

## Data analysis

### Exclusions

Final group sizes are given for each experiment above. Four fiber photometry rats were excluded (10-s delay cohort: 1 rat with poor GCaMP expression; 5-s delay cohort: 2 rats failed to perform task, 1 rat for poor photometry data quality).

### Statistical analysis

Analyses were conducted in Python 3.9.12 using pandas 2.3.3, NumPy 1.22.4, SciPy 1.7.3, statsmodels 0.13.2, and pingouin 0.5.5. All tests were two-tailed and alpha was set at p < 0.05. Factorial designs were analyzed by repeated-measures or mixed ANOVA. Each experiment was evaluated on a single pre-specified omnibus term (devaluation, state × cue, or drug × cue); planned comparisons on that term were reported as paired or independent t tests, as appropriate. Effect sizes are reported as partial η² for ANOVA terms and Cohen’s d (or dz for within-subject comparisons) for t tests, each with a 95% confidence interval obtained by inverting the corresponding noncentral distribution: the noncentral F for partial η² and the noncentral t for d and dz. Subject means were residualized on counterbalancing assignment before analysis and plotting. Trial-level relationships between photometry and behavior were modeled by generalized estimating equations, and CS-evoked transients were evaluated by hierarchical bootstrap, both as described below.

### Behavioral analyses

Mean lever press rates (responses per min) for the last 3 sessions of initial instrumental training were used to assess baseline performance. Baseline conditioned food-port activity during the last 3 Pavlovian training sessions was assessed by computing the difference in responding during CS periods relative to pre-CS baseline periods [Δ = CS – Pre-CS; 10-s each], calculated separately for each CS type. Difference scores were also used to calculate cue-induced changes in press rate and food-port activity during PIT tests. Because lever pressing and food-port entries are commonly performed together as instrumental response sequences, we focused our analyses on the frequency of spontaneous (or press-independent) bouts of food-port activity, which were defined as entries that occurred at least 2.5-s after the last detected food-port entry or lever press, as in earlier studies (Marshall and Ostlund, 2018; Marshall et al., 2020; Halbout et al., 2022; Malvaez et al., 2026). Our primary analysis of results from the hunger-to-satiety shift study focused on data from the first half of each test (trials 1-5) to minimize the disruptive response-suppressive effects of satiety and extinction (repeated testing) on later trials. Full-session results are provided in Figure S2 and generally align with our primary analysis.

Prior research with the PIT task used here found that rats show more vigorous lever pressing during cues signaling a low-probability of reward (CS_30_) than cues signaling a high probability of reward (CS_100_) (Marshall et al., 2020; Malvaez et al., 2026). We propose that this cue-specificity reflects the ability to exert top-down control over instrumental reward-seeking when imminent reward delivery is strongly predicted. We quantified this as a cue-specificity score (Δ press for CS_30_ – Δ press for CS_100_). We performed supplementary analyses to characterize response competition between lever pressing and food-port occupancy during PIT task performance. This included analysis of pooled data from experiments involving both CS_30_ and CS_100_ (and limited to tests conducted under control conditions when typical performance should be expected, which included the hungry test in the satiety-shift experiment, the photometry experiment, and vehicle tests (both hM4Di and GFP groups) from the chemogenetic inhibition experiment) (Figure 5a–e). We also computed an opportunity-adjusted press rate, to determine whether cue-specific differences in pressing could be produced by cue-specific differences in the time available to press. This measure expresses presses per second of cue time not spent occupying the food port, and was computed separately for each subject and cue: presses and available time [max(CS duration – port-occupancy seconds, 0.1 s)] were summed across that cue’s trials before dividing, so that trials are weighted by the time available in them; the same adjustment was applied to the pre-CS period before subtraction. This adjustment assumes that time spent in the food port would otherwise have been spent pressing, at the rate achieved during the remainder of the cue period. Effects of interest are reported both unadjusted and adjusted (Tables 1–4).

### Fiber photometry

Photometry signals were extracted and downsampled to 16.15 Hz. Ca^2+^-dependent (465 nm) and isosbestic (405 nm) signals were high-pass filtered (90 s period, 0.0111 Hz cutoff) and low-pass filtered (5 Hz cutoff). Robust regression (Huber regression, ε = 1.35), similar to (Keevers and Jean-Richard-Dit-Bressel, 2025), was used to fit the isosbestic signal onto the Ca^2+^-dependent signal, and ΔF/F was calculated as (Ca^2+^-dependent signal − fitted 405 nm) / fitted 405 nm. Outlier ΔF/F values exceeding 7 SD from the session mean were then identified in the continuous photometry signal and replaced via linear interpolation (<0.5% of samples from any session). Photometry signals were extracted for CS presentations (onset at t = 0) using three analysis windows: (1) full CS epoch (−3 to 13 s), z-scored to the pre-CS baseline (−1 to 0 s); (2) CS onset (−1 to 3 s), using full CS normalization; (3) CS offset (9 to 13 s), re-normalized to a 1 s baseline immediately preceding offset (9 to 10 s). Trials were further categorized based on whether lever pressing occurred during the CS period (0– 10 s).

We used a hierarchical bootstrapping approach to determine statistically significant CS-evoked transients (Jean-Richard-dit-Bressel et al., 2020; Tan et al., 2026). Each session contributed one mean trace, and 10,000 bootstrap iterations resampled sessions (with replacement) to generate a distribution of group means. A 95% confidence interval was calculated using the 2.5th and 97.5th percentiles, then expanded by a factor of √(n / (n − 1)) to correct for narrowness bias. Significant transients were defined as periods where the 95% CI did not contain 0% ΔF/F for at least 0.33 s (consecutive threshold). Temporal smoothing (3 timepoints) was applied to bridge brief gaps (≤ 186 ms) in significance. This approach was applied to: (1) CS onset and offset epochs for CS_100_ and CS_30_ separately, plus CS_100_ − CS_30_ contrasts; (2) full CS trials with vs. without lever pressing for each CS type, plus press − no press contrasts; (3) full CS_100_ vs. CS_30_ for descriptive comparison.

To examine the relationship between trial-by-trial dmPFC GCaMP activity and lever pressing, we used generalized estimating equations (GEE) with a binomial family and logit link function to model binary pressing outcomes (press vs. no press during CS presentations). Mean GCaMP activity during the CS period (0–10 s) was z-scored and entered as a predictor, along with CS type (CS_100_ vs. CS_30_) and their interaction. Trial order (1–10, mean-centered) was included as a covariate to control for within-session temporal effects (e.g., extinction). An exchangeable correlation structure accounted for within-session clustering of trials. Separate models were fit for each CS type to derive simple slopes, and a full interaction model tested whether the GCaMP-pressing relationship was CS-dependent.

### Code accessibility

All photometry and behavioral analyses were performed using custom Python code. The analysis code and the processed data required to reproduce every statistic and figure reported here are available from the corresponding author on request.

## Results

We began by asking the extent to which the regulatory effects of cues that signal imminent reward delivery on instrumental reward seeking are goal-directed. It was recently shown that this regulatory influence is attenuated, or even reversed, when cues predict relatively small or devalued rewards (Marshall et al., 2023). We hypothesized that this reflects the use of a top-down flexible control process in which cues for imminent reward flexibly constrain the motivational impulse to engage in instrumental reward seeking to promote behaviors that are normally more advantageous in such contexts, specifically approaching and waiting for the predicted reward delivery (Ostlund and Marshall, 2021). According to this view, motivation for instrumental reward seeking should not be suppressed in response to a cue that predicts imminent delivery of a low or devalued reward. We tested this hypothesis using a variant of the PIT task that is optimized to assay the regulatory influence of reward predictive cues on instrumental reward seeking.

### Reward devaluation releases cue-evoked instrumental reward seeking

Briefly, rats were given Pavlovian delay conditioning in which two 10-s auditory cues (clicker or tone) signaled delivery of different food outcomes (grain pellet or SCM solution) at cue offset, on either 100% or 30% of trials (between-subjects) (Figure 1a). Baseline conditioned food-port activity (CS − PreCS) did not differ across the two cues prior to devaluation (Figure S1). Rats were then trained to lever press for a distinct banana pellet reward and established vigorous baseline press rates (Figure S1). This switch in reward identity across training phases allowed us to then change the relative value of Pavlovian rewards without impacting the value of the instrumental reward, thereby exposing the influence of reward value on the general or nonspecific motivational properties of the CS (Corbit and Balleine, 2015). After training, one of the two Pavlovian rewards was selectively devalued through conditioned taste aversion training, leading rats to reject it during daily consumption opportunities while intake of the still-valued outcome remained at ceiling (Figure 1b).

**Figure 1.**
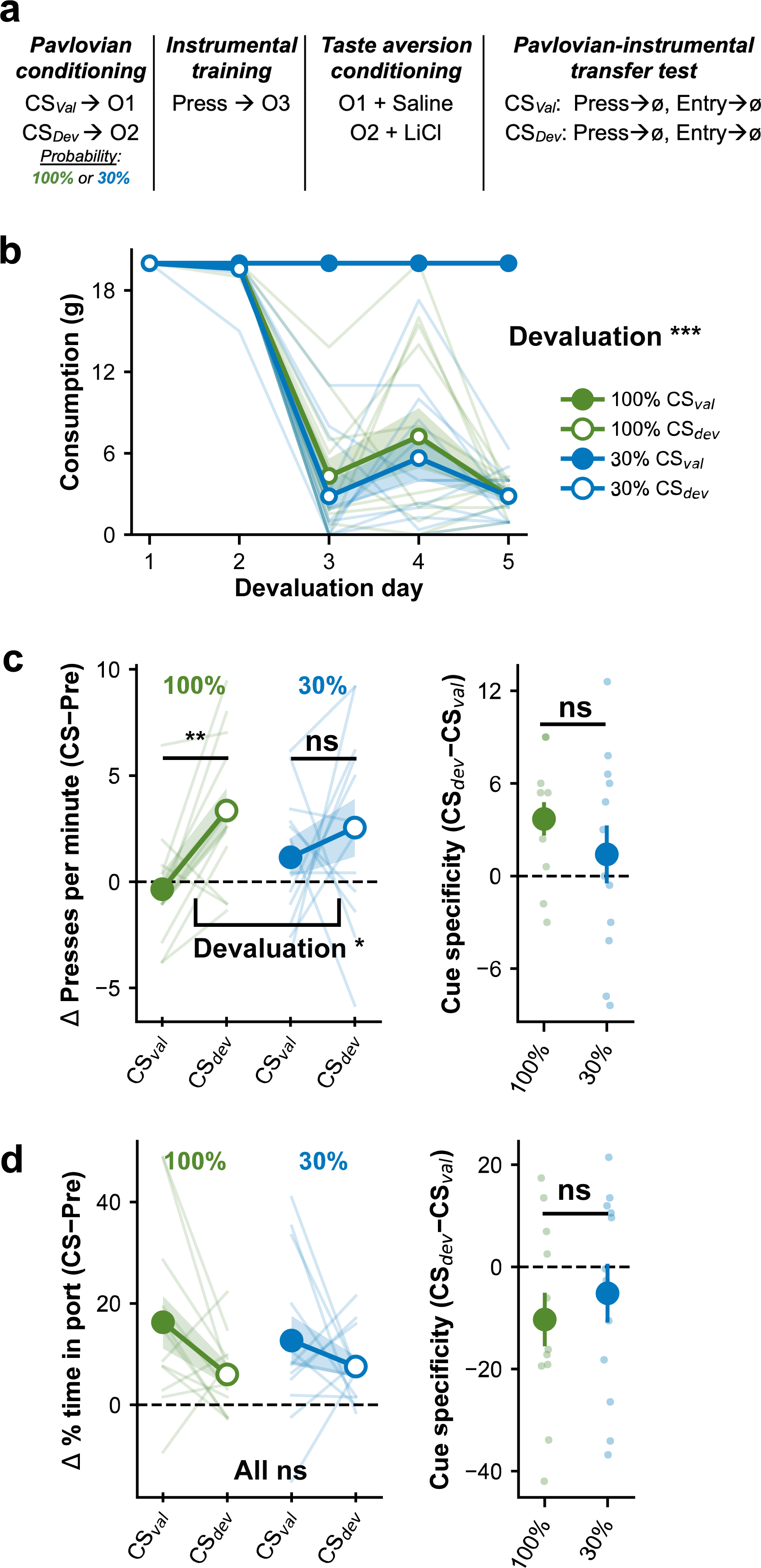
Devaluing the predicted reward releases cue-evoked instrumental reward seeking. ***a***, Task schematic. CS*_Val_* and CS*_Dev_*, 10-s auditory conditioned stimuli (click or tone), each coterminating with a distinct food outcome (O1, O2) on 100% or 30% of trials (between-subjects). Press, lever press earns a third, distinct food outcome (O3). Taste aversion conditioning, one outcome paired with LiCl and the other with saline, defining CS*_Dev_* and CS*_Val_* for each rat. At the transfer test, both cues are presented while lever pressing and food-port activity are measured. Ø, no rewards were delivered at test. ***b***, Consumption of the CS*_Val_*- and CS*_Dev_*-predicted outcomes across the five taste-aversion conditioning sessions, plotted separately for the 100% and 30% groups. Condition × Day RM-ANOVA, condition, F(1,23) = 675.46; p < 0.0001; condition × day, F(4,92) = 167.36; p < 0.0001; day, F(4,92) = 167.36; p < 0.0001. Consumption of the valued outcome was invariant across sessions, so the day and condition × day terms are numerically identical. Day 5, CS*_Val_*- versus CS*_Dev_*-predicted outcome, t(23) = 55.67; p < 0.0001. ***c***, Left, Cue-evoked change (Δ) in lever pressing from the preceding baseline period during CS*_Val_* and CS*_Dev_*, by reward-probability group. Mixed ANOVA, devaluation × group, F(1,22) = 1.13; p = 0.30; devaluation, F(1,22) = 5.54; p = 0.028; group, F(1,22) = 0.12; p = 0.73. Planned comparisons, 100%, CS*_Val_* versus CS*_Dev_*, t(11) = −3.43; p = 0.0057; 30%, CS*_Val_* versus CS*_Dev_*, t(11) = −0.75; p = 0.47. Right, Cue-specificity score (Δ presses during CS*_Dev_* − Δ presses during CS*_Val_*; positive values indicate greater pressing during CS*_Dev_*) versus zero, 100%, t(11) = 3.43; p = 0.0057; 30%, t(11) = 0.75; p = 0.47; 100% versus 30%, t(22) = 1.06; p = 0.30. ***d***, Left, Cue-evoked change in percentage of time spent in the food port from the preceding baseline period. Mixed ANOVA, devaluation × group, F(1,22) = 0.44; p = 0.52; devaluation, F(1,22) = 3.96; p = 0.059; group, F(1,22) = 0.07; p = 0.79. Planned comparisons, 100%, CS*_Val_* versus CS*_Dev_*, t(11) = 1.97; p = 0.075; 30%, CS*_Val_* versus CS*_Dev_*, t(11) = 0.90; p = 0.39. Right, Cue-specificity score (Δ % time in port during CS*_Dev_* − CS*_Val_*) versus zero, 100%, t(11) = −1.97; p = 0.075; 30%, t(11) = −0.90; p = 0.39; 100% versus 30%, t(22) = −0.66; p = 0.52. n = 24 rats (12 per group). Data presented as mean ± SEM; subject means residualized on counterbalancing group. Baseline and CS analysis windows, 10 s. Planned comparisons protected by the omnibus devaluation term. *p < 0.05; **p < 0.01; ***p < 0.001.

Rats were then given a PIT test, during which the two cues were intermittently presented to assess their influence on instrumental reward seeking. Devaluing the predicted outcome released cue-evoked instrumental reward seeking (Figure 1c; devaluation, F(1,22) = 5.537, p = 0.028). Rats pressed at a higher rate during the cue that signaled the devalued outcome (CS_Dev_) than during the cue that signaled the still-valued outcome (CS_Val_). The effect was carried by the group trained with cues that predicted reward on 100% of trials (t(11) = −3.43, p = 0.0057); the same comparison was not reliable in the 30% group (t(11) = −0.75, p = 0.47). Devaluation had no reliable effect on cue-evoked food-port occupancy time in either group (Figure 1d). Rats in the 100% group did, however, initiate fewer bouts of food-port entry when the devalued reward was predicted (F(1,11) = 5.42, p = 0.040), an effect that was not reliable in the 30% group (Figure S1). Removing the value of the predicted reward therefore unmasked the cue’s latent capacity to invigorate instrumental reward seeking, while leaving the duration of conditioned goal-approach unchanged.

### A shift from hunger to satiety abolishes cue-specific regulation of instrumental reward seeking

In a new group of rats, as a converging test of the role of reward value on task performance, we assessed whether a shift in homeostatic need state, from hunger to satiety, alters the behavioral effects of cues signaling imminent reward. For this and all subsequent experiments reported here, rats were given Pavlovian conditioning with two cues that were probabilistically reinforced to signal either a 100% (CS_100_) or 30% (CS_30_) chance of imminent reward delivery (within-subject; Figure 2a), which supported differential levels of conditioned food-port activity by the end of training (Figure S2). Rats were then trained to lever press for food reward (Figure S2), after which they were administered a pair of PIT tests conducted while subjects were Hungry or Sated. This manipulation altered how the two cues influenced instrumental reward seeking (Figure 2b; state × CS, F(1,39) = 5.442, p = 0.025). Consistent with prior research (Marshall et al., 2020; Malvaez et al., 2026), hungry rats pressed at a higher rate during CS_30_ than during CS_100_, withholding instrumental responding to the cue that strongly predicted imminent reward delivery. Under satiety, the two cues elicited comparable increases in pressing, abolishing this cue-specific regulation. Satiety also lowered the overall rate of lever pressing measured during the pre-CS baseline period, indicating that the loss of cue-specific regulation was accompanied by a general downward shift in the instrumental behavior on which the cues act, rather than by a selective disruption of the regulatory process itself (Figure S2). Conditioned food-port activity, in contrast, was unaffected by this manipulation, such that both cues continued to elicit robust increases in food-port occupancy, and the selective preference for CS_100_ over CS_30_ was preserved in both states (Figures 2c and S2).

**Figure 2.**
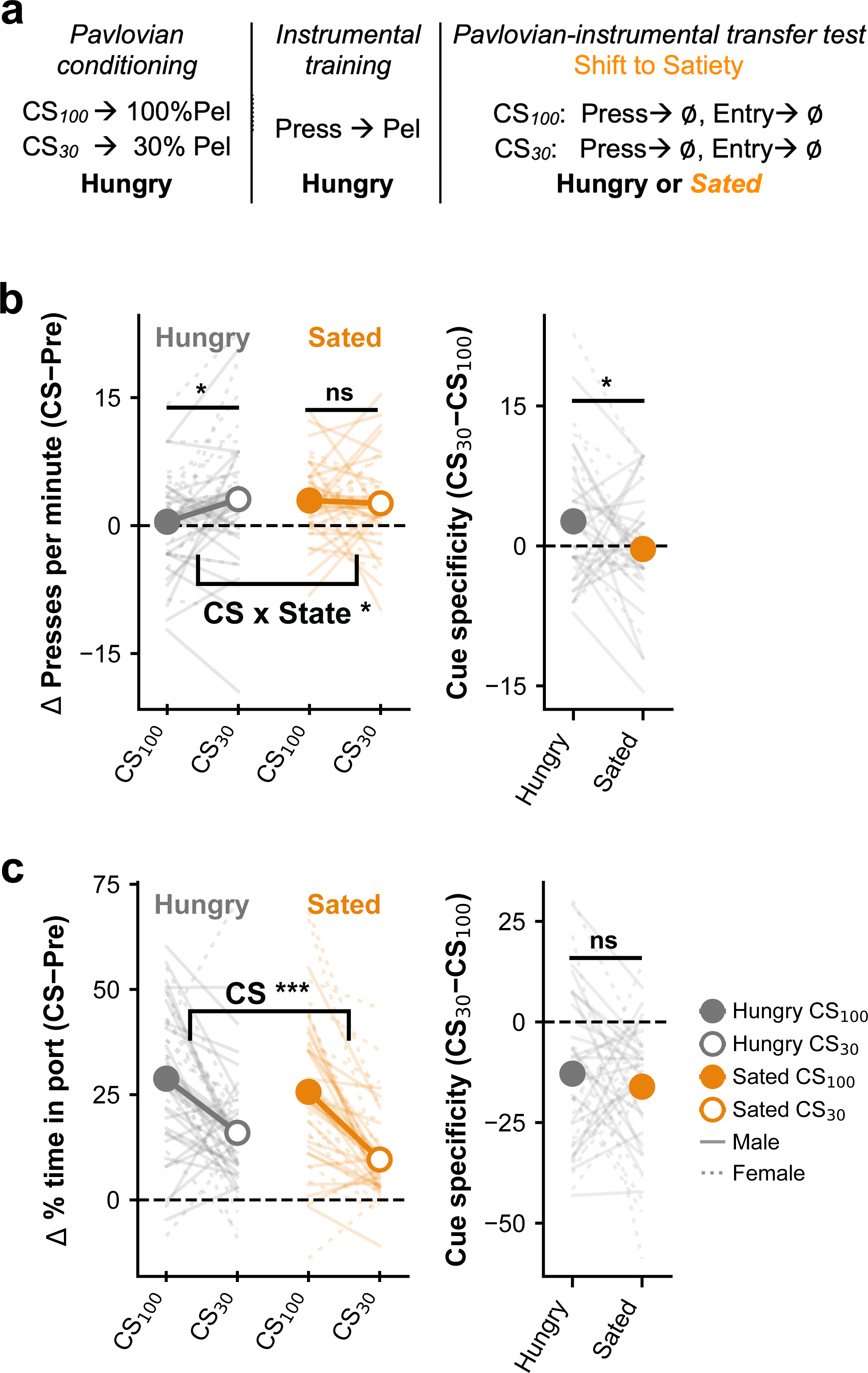
Satiety abolishes cue-specific regulation of instrumental reward seeking while leaving cue-directed food-port checking intact. ***a***, Task schematic. CS_100_ and CS_30_, 10-s auditory conditioned stimuli (click or tone) coterminating with a food-pellet (Pel) reward on 100% or 30% of trials, counterbalanced. Press, lever press earns the same food-pellet reward. Ø, no rewards were delivered at test. Rats were trained hungry and received transfer tests in both the hungry and sated states in a within-subject crossover. ***b***, Left, Cue-evoked change (Δ) in lever pressing from the preceding baseline period during CS_100_ and CS_30_, in the hungry and sated states. RM-ANOVA, state × CS, F(1,39) = 5.44; p = 0.025; CS, F(1,39) = 2.14; p = 0.15; state, F(1,39) = 1.08; p = 0.31. Planned comparisons, hungry, CS_100_ versus CS_30_, t(39) = −2.44; p = 0.019; sated, CS_100_ versus CS_30_, t(39) = 0.35; p = 0.73. Right, Cue- specificity score (Δ presses during CS_30_ − Δ presses during CS_100_; positive values indicate greater pressing during CS_30_) versus zero, hungry, t(39) = 2.44; p = 0.019; sated, t(39) = −0.35; p = 0.73; hungry versus sated, t(39) = 2.33; p = 0.025. ***c***, Left, Cue- evoked change in percentage of time spent in the food port from the preceding baseline period. RM-ANOVA, state × CS, F(1,39) = 0.67; p = 0.42; CS, F(1,39) = 53.84; p < 0.0001; state, F(1,39) = 3.30; p = 0.077. Planned comparisons, hungry, CS_100_ versus CS_30_, t(39) = 4.26; p < 0.001; sated, CS_100_ versus CS_30_, t(39) = 6.33; p < 0.0001. Right, Cue-specificity score (Δ % time in port during CS_30_ − CS_100_) versus zero, hungry, t(39) = −4.26; p < 0.001; sated, t(39) = −6.33; p < 0.0001; hungry versus sated, t(39) = 0.82; p = 0.42. Satiety reduced overall pressing: pre-CS response rate, hungry versus sated, t(39) = 3.68; p = 0.001. n = 40 rats (24 male, 16 female). Test data are restricted to the first five presentations of each cue (see Materials and Methods and Supplemental Material for full session results). Data presented as mean ± SEM; subject means residualized on counterbalancing group. Baseline and CS analysis windows, 10 s. Filled symbols, CS_100_; open symbols, CS_30_. Males, solid lines; females, dotted lines. Planned comparisons protected by the omnibus state × CS term. *p < 0.05; ***p < 0.001.

### dmPFC activity scales with cued reward probability and signals reward omission

These findings show that the regulatory influence of cues that signal imminent reward is flexible, depending both on the value of the predicted reward and on the animal’s motivational state. Devaluing the predicted reward reversed the direction of the cue’s effect on pressing rather than simply removing it, exposing a latent motivational influence that is otherwise held in check when imminent reward is expected. This argues for an active, expectancy-dependent control process rather than a simple absence of motivation, and raises the question of what circuitry implements it. To investigate the role of the dmPFC in this process, we used fiber photometry to determine how dmPFC activity is modulated by such cues and their influence on instrumental reward seeking. If the dmPFC uses cue-elicited reward predictions to regulate reward seeking, its activity should distinguish cues that differ in predicted reward probability, and should also track whether that regulation is expressed on a given trial. Two patterns would satisfy the latter prediction, with opposite interpretations. If dmPFC activity indexes the implementation of control, it should be greatest on trials where reward seeking is successfully withheld. If instead it indexes the demand for control (i.e., the strength of the impulse to be overcome), it should be greatest on the trials where that impulse prevails and the rat presses. We therefore examined cue-evoked activity separately for trials with and without a lever press.

Rats were given intracranial AAV injections to express the calcium indicator GCaMP in dmPFC neurons (Figure 3a). They were then given probabilistic Pavlovian conditioning (Figure 3b), developing higher levels of conditioned food-port activity to CS_100_ than CS_30_ (Figure S3). After instrumental training to establish lever pressing for food reward (Figure S3), rats were given a PIT test while dmPFC GCaMP activity was recorded. In this cohort, the influence of CS presentations on lever pressing (Figure 3c) and food-port occupancy (Figure 3d) was variable across subjects and did not reliably depend on predicted reward probability, possibly reflecting the constraint of the patch cable or a generalization decrement across training and testing chambers. New bouts of food-port activity were nonetheless more frequent during CS_100_ than CS_30_ (Figure 3e), confirming that rats discriminated between the two cues at test. dmPFC neuron bulk GCaMP activity was strongly and differentially modulated by these cue presentations (Figure 3f,g). A pronounced elevation in GCaMP occurred within the first few seconds of CS onset and was significantly larger for CS_100_ than CS_30_, likely reflecting predicted reward probability. GCaMP activity also showed a transient dip shortly after CS termination, the expected delivery time for the omitted reward.

**Figure 3.**
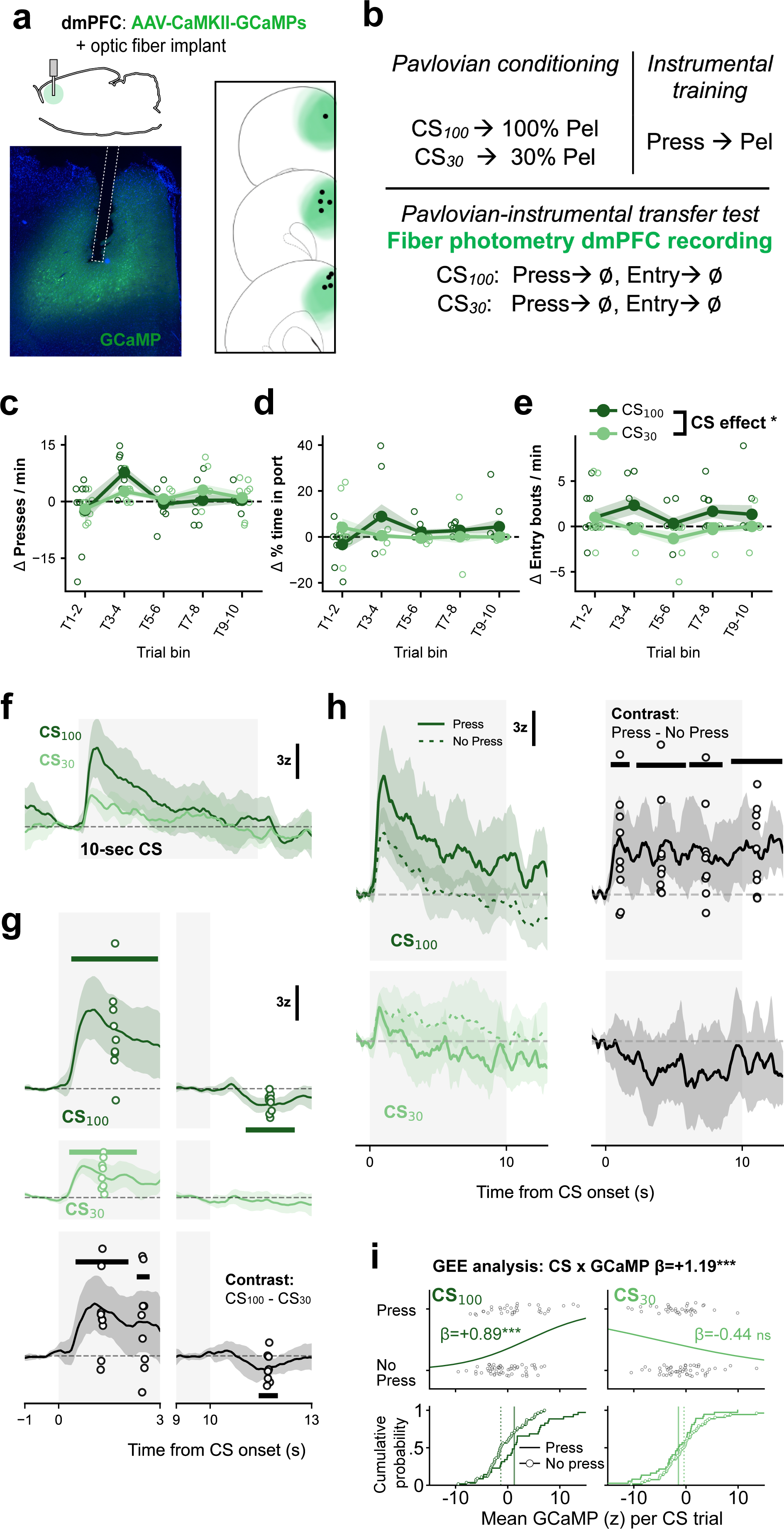
dmPFC population activity scales with cued reward probability and signals the omission of a strongly predicted reward. ***a***, Fiber photometry approach for imaging GCaMP6s fluorescence changes in dmPFC neurons (AAV-CaMKII-GCaMP6s and optical fiber implant). ***b***, Task training and test schematic. CS_100_ and CS_30_, 10-s auditory conditioned stimuli (click or tone) coterminating with a food-pellet (Pel) reward on 100% or 30% of trials, counterbalanced. Press, lever press earns the same food-pellet reward. Ø, no rewards were delivered at test. Photometry was recorded throughout the transfer test. ***c***, Cue-evoked change (Δ) in lever pressing from the preceding baseline period, in five successive 2-trial bins. RM- ANOVA, CS × trial bin, F(4,32) = 1.61; p = 0.22; CS, F(1,8) = 0.00; p = 1.00; trial bin, F(4,32) = 6.29; p = 0.0029. ***d***, Cue-evoked change in percentage of time spent in the food port. RM-ANOVA, CS × trial bin, F(4,32) = 2.68; p = 0.097; CS, F(1,8) = 0.78; p = 0.40; trial bin, F(4,32) = 0.93; p = 0.41. ***e***, Cue-evoked change in food-port entry bouts. RM-ANOVA, CS × trial bin, F(4,32) = 1.01; p = 0.39; CS, F(1,8) = 9.00; p = 0.017; trial bin, F(4,32) = 1.28; p = 0.30. ***f***, Mean GCaMP6s fluorescence (z-score) during CS_100_ and CS_30_; bar, 10-s CS. ***g,*** *GCaMP6s fluorescence (z-score) in the CS-onset and CS- offset epochs, aligned to CS onset.* Bootstrap, 10,000 iterations, narrowness-corrected 95% CI, 0.33-s consecutive-significance threshold. CS onset (−1 to 3 s), CS_100_, elevated 0.38–2.94 s; CS_30_, elevated 0.31–2.31 s; CS_100_ − CS_30_, 0.50–2.69 s. CS offset (9–13 s, renormalized to a 9–10-s baseline), CS_100_, decreased 11.06–12.50 s; CS_30_, no significant window; CS_100_ − CS_30_, 11.50–12.00 s. ***h***, Mean GCaMP6s fluorescence on trials with and without a lever press during the cue, and the press minus no-press contrast. CS_100_, press exceeded no-press at 0.38–1.75 s, 2.31–5.88 s, 6.12–8.56 s, and 9.19–12.94 s; CS_30_, no significant window; CS × behavior interaction, 0.94–1.56 s, 2.62–5.19 s, 5.75–8.12 s, 8.38–9.25 s, 10.12–10.88 s, and 11.31–12.94 s. ***i***, Top, Trial- level logistic generalized estimating equation predicting whether a lever press occurred during the cue from cue-evoked GCaMP (exchangeable working correlation, clustered by rat; 9 rats, all 10 trials per cue, GCaMP standardized per SD, trial order included as a covariate). CS × GCaMP, β = +1.19; p < 0.0001. Simple slopes from per-cue fits of the same model, CS_100_, β = +0.89; p < 0.001; CS_30_, β = −0.44; p = 0.079. Bottom, Cumulative distribution of mean cue-evoked GCaMP across trials on which a press did and did not occur, by cue. n = 9 rats (all male). Behavioral panels, mean ± SEM; photometry traces, group mean with narrowness-corrected bootstrap 95% confidence interval. Trial-bin ANOVAs, Greenhouse–Geisser corrected. *p < 0.05; **p < 0.01; ***p < 0.001.

Consistent with a negative reward prediction error, this reduction differed across cues and was reliable for the highly predictive CS_100_ but not the weakly predictive CS_30_ (Figure 3g). This offset response appeared to track the trained time of reward delivery rather than cue offset per se. In a separate group of rats trained with a 5-s delay, dmPFC activity dipped below baseline after that group’s expected time of reward delivery, with no further modulation at CS offset (Figure S3m).

### Cue-evoked dmPFC activity predicts instrumental responding on a trial-by-trial basis

To examine the behavioral relevance of this CS-elicited dmPFC GCaMP activity, we took advantage of the trial-to-trial variability in task performance and separated trials based on whether rats engaged in lever pressing (binary: press or no press) during the CS period. CS_100_ evoked larger GCaMP responses on trials when that cue elicited lever pressing than on trials when it did not. No comparable difference was detected for CS_30_ (Figure 3h). Because this cohort showed no group-level difference in cue-evoked pressing, trial-level analysis provides the more informative test of whether dmPFC activity tracks instrumental performance. We therefore examined these relationships at the level of individual trials using logistic generalized estimating equations (GEE) to model cue-evoked lever press behavior while accounting for within-session clustering of trials. This analysis confirmed a cue-dependent relationship between dmPFC GCaMP activity and lever pressing (Figure 3i; GCaMP × CS type interaction: β = +1.193, p < 0.0001). Higher GCaMP activity during CS_100_ was associated with an increased likelihood of pressing, whereas during CS_30_ the relationship showed a marginal trend in the opposite direction (β = −0.44, p = 0.079). These results indicate that the dmPFC is activated by reward-predictive cues, encodes information about upcoming reward probability, and is associated with lever-press performance, particularly when imminent reward is predicted, signaling a demand for greater control over such behavior.

### Chemogenetic dmPFC inhibition abolishes cue-specific regulation of instrumental reward seeking

We next used a chemogenetic strategy to investigate whether the dmPFC is causally involved in using cue-elicited reward predictions to regulate instrumental reward seeking. Given prior evidence that stimulating this region suppresses cue-evoked instrumental performance (Halbout et al., 2022), together with our finding that cue-evoked dmPFC activity scales with predicted reward probability and tracks trial-by-trial pressing, we asked whether inhibiting it would instead impair the ability to use reward predictions to negatively regulate that behavior. Rats underwent surgery to virally express the inhibitory designer receptor hM4Di in the dmPFC (Figure 4a). After training (Figure 4b; Figure S4), rats were administered PIT tests following pretreatment with the hM4Di agonist CNO or vehicle. In hM4Di-expressing rats, CNO abolished the cue-specific regulation of lever pressing (Figure 4c; drug × CS, F(1,13) = 15.113, p = 0.0019). Under vehicle these rats withheld pressing during CS_100_ relative to CS_30_, whereas under CNO the two cues elicited comparable levels of pressing. A separately run GFP control group, analyzed independently, showed cue-specific regulation of lever pressing that was intact and unaffected by drug treatment (Figure 4e). dmPFC inhibition had a distinct effect on conditioned food-port activity. In hM4Di rats, CNO reduced the overall time spent in the food port during cue presentations, an effect that was clearest during CS_100_. The cue-specificity of this behavior was preserved, however, with CS_100_ continuing to elicit more food-port activity than CS_30_ under both drug conditions, although the drug × CS interaction approached significance (F(1,13) = 4.05, p = 0.065; Figure 4d; Figure S4). GFP control rats likewise showed intact, drug-independent cue-directed food-port activity (Figure 4f). The disruptive effect of dmPFC inhibition was therefore limited to the use of reward-predictive cues to flexibly regulate instrumental reward seeking.

**Figure 4.**
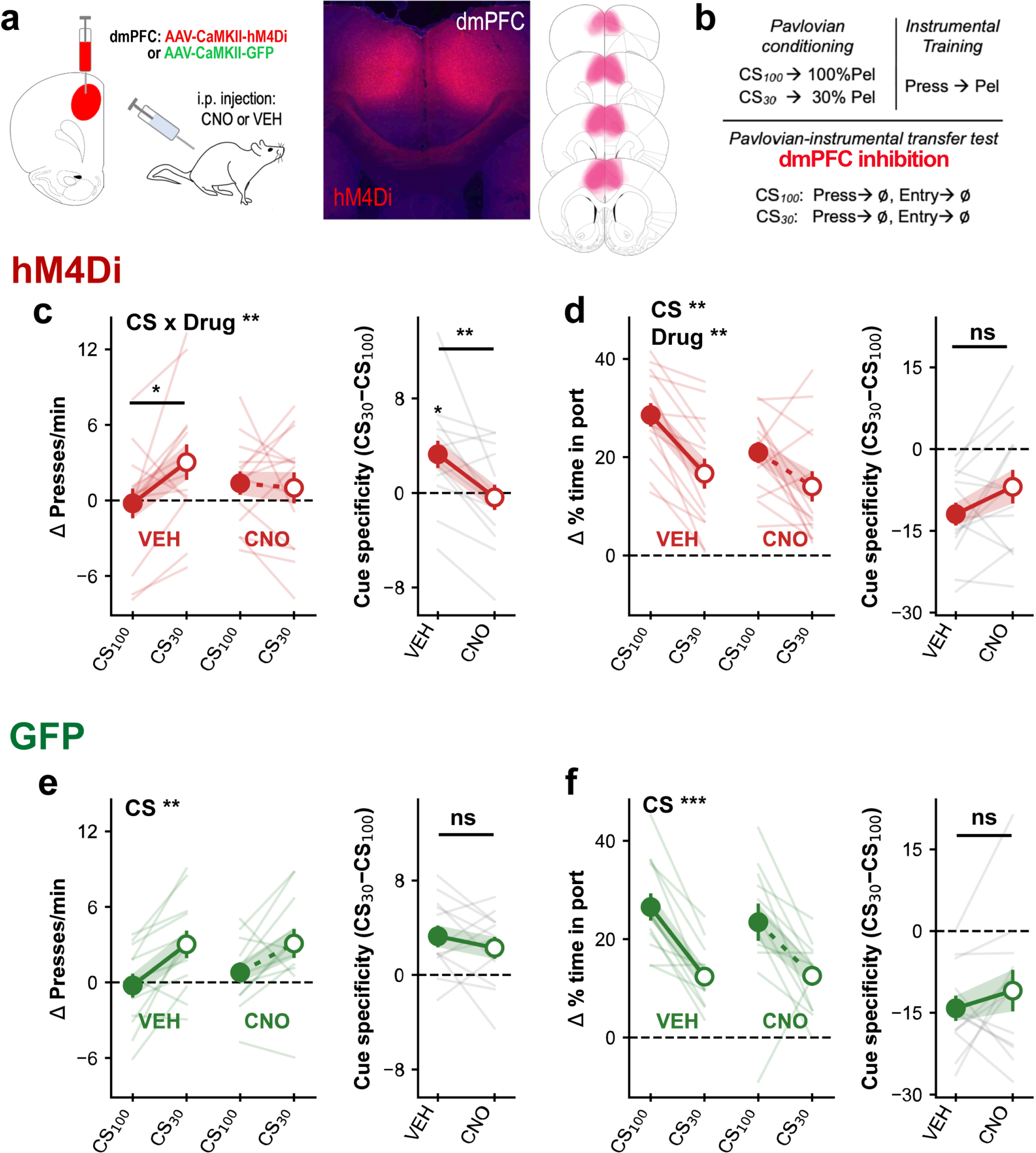
Chemogenetic inhibition of dmPFC abolishes cue-specific regulation of instrumental reward seeking while leaving cue-directed food-port checking intact. ***a***, Chemogenetic approach. Rats received bilateral dmPFC infusions of AAV-CaMKII- hM4Di or AAV-CaMKII-GFP and were pretreated with CNO or vehicle (i.p.) before each transfer test in a within-subject crossover. Middle, Representative immunofluorescence image of dmPFC hM4Di-mCherry expression. Right, hM4Di expression map. ***b***, Task training and test schematic. CS_100_ and CS_30_, 10-s auditory conditioned stimuli (click or tone) coterminating with a food-pellet (Pel) reward on 100% or 30% of trials, counterbalanced. Press, lever press earns the same food-pellet reward. Ø, no rewards were delivered at test. ***c***, Left, Cue-evoked change (Δ) in lever pressing from the preceding baseline period during CS_100_ and CS_30_, under vehicle and CNO, in hM4Di- expressing rats. RM-ANOVA, drug × CS, F(1,13) = 15.11; p = 0.0019; CS, F(1,13) = 2.09; p = 0.17; drug, F(1,13) = 0.04; p = 0.84. Planned comparisons, vehicle, CS_100_ versus CS_30_, t(13) = −2.85; p = 0.014; CNO, CS_100_ versus CS_30_, t(13) = 0.34; p = 0.74. Right, Cue-specificity score (Δ presses during CS_30_ − Δ presses during CS_100_) versus zero, algebraically equivalent to the paired comparisons at left: vehicle, t(13) = 2.85; p = 0.014; CNO, t(13) = −0.34; p = 0.74; vehicle versus CNO is the drug × CS term above. CNO did not shift pressing to either cue on its own, CS_100_, t(13) = −1.34; p = 0.20; CS_30_, t(13) = 1.87; p = 0.085. ***d***, Left, Cue-evoked change in percentage of time spent in the food port, hM4Di group. RM-ANOVA, drug × CS, F(1,13) = 4.05; p = 0.065; CS, F(1,13) = 16.38; p = 0.0014; drug, F(1,13) = 11.72; p = 0.0045. Planned comparisons, vehicle, CS_100_ versus CS_30_, t(13) = 5.72; p < 0.0001; CNO, CS_100_ versus CS_30_, t(13) = 2.23; p = 0.044. CNO reduced food-port time during CS_100_, vehicle versus CNO, t(13) = 3.34; p = 0.0053, but not during CS_30_, t(13) = 1.69; p = 0.11. Right, Cue-specificity score (Δ % time in port during CS_30_ − CS_100_) versus zero, vehicle, t(13) = −5.72; p < 0.0001; CNO, t(13) = −2.23; p = 0.044; vehicle versus CNO is the drug × CS term above. ***e***, Left, Cue- evoked change in lever pressing in GFP control rats. RM-ANOVA, drug × CS, F(1,11) = 1.04; p = 0.33; CS, F(1,11) = 12.54; p = 0.0046; drug, F(1,11) = 1.15; p = 0.31. Planned comparisons, vehicle, CS_100_ versus CS_30_, t(11) = −3.58; p = 0.0043; CNO, CS_10_0 versus CS_30_, t(11) = −2.48; p = 0.031. Right, Cue-specificity score versus zero, vehicle, t(11) = 3.58; p = 0.0043; CNO, t(11) = 2.48; p = 0.031; vehicle versus CNO is the drug × CS term above. ***f***, Left, Cue-evoked change in percentage of time spent in the food port, GFP group. RM-ANOVA, drug × CS, F(1,11) = 1.00; p = 0.34; CS, F(1,11) = 21.51; p = 0.0007; drug, F(1,11) = 0.65; p = 0.44. Planned comparisons, vehicle, CS_100_ versus CS_30_, t(11) = 6.13; p < 0.0001; CNO, CS_10_0 versus CS_30_, t(11) = 2.86; p = 0.016. Right, Cue-specificity score versus zero, vehicle, t(11) = −6.13; p < 0.0001; CNO, t(11) = −2.86; p = 0.016; vehicle versus CNO is the drug × CS term above. hM4Di, n = 14 rats; GFP, n = 12 rats. The two viruses were run as separate cohorts and are analyzed independently throughout. Data presented as mean ± SEM; subject means residualized on counterbalancing group. Baseline and CS analysis windows, 10 s. Filled symbols, CS_100_; open symbols, CS_30_. Solid lines, vehicle; dashed lines, CNO. Planned comparisons protected by the omnibus drug × CS term. *p < 0.05; **p < 0.01; ***p < 0.001.

### Response competition is local and does not account for cue-specific regulation of instrumental behavior

Cues in this task elicit two incompatible responses: approach to the food port and pressing on the lever. Because rats occupied the food port more during CS_100_ than during CS_30_, the lower press rate during CS_100_ could in principle reflect the distribution of the animal’s time rather than any top-down regulatory influence over instrumental performance.

Competition of this kind can constrain pressing only if port occupancy consumes enough of the cue period to crowd out presses the rat would otherwise emit, a condition that was rarely met. Across control-condition tests (see Behavioral Analyses for details), port occupancy accounted for 3.16 ± 0.20 s of the 10-s CS_100_ and 1.90 ± 0.17 s of CS_30_ (31.6 ± 2.0% and 19.0 ± 1.7% of the cue period), leaving 6.84 ± 0.20 s and 8.10 ± 0.17 s available for pressing, during which rats emitted only 0.89 ± 0.07 and 1.26 ± 0.11 presses per trial, respectively. Press rate did decline as occupancy increased, but this relationship was confined to the small minority of observations in which occupancy was high (Figure 5a–e; see also Figure S5 for the reward devaluation experiment, which is not part of this pooled sample). Rats thus had ample opportunity to press during both cues, leaving little scope for competition over time to generate a systematic difference between them.

**Figure 5.**
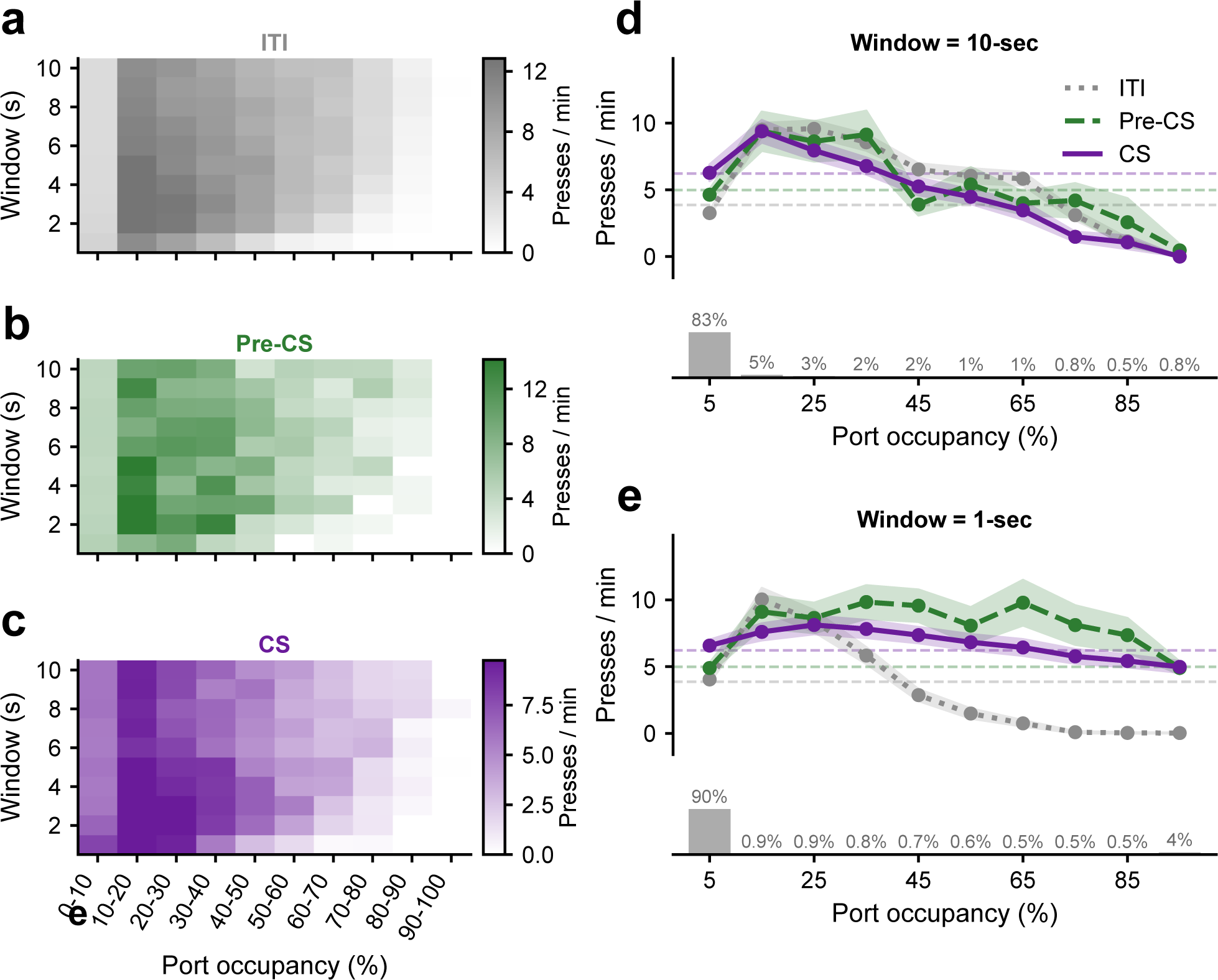
Competition with food-port occupancy is real but rare, and does not account for cue-specific regulation of instrumental performance. Lever pressing and food-port approach cannot be performed at the same moment, and cues signaling imminent reward draw rats to the port. Cue-specific differences in press rate could therefore reflect how an animal distributed its time rather than regulation of instrumental performance. Competition can constrain pressing only if port occupancy, the proportion of the cue period a rat spends inside the food port, is high enough to displace presses that rat would otherwise have emitted. Panels ***a–e*** characterize how much of the cue period was occupied and how press rate varied with occupancy in the pooled control-condition sample (N = 75 tests): the hungry test from the satiety-shift experiment (n = 40), the photometry experiment (n = 9), and the vehicle tests of both virus groups from the chemogenetic inhibition experiment (n = 26). The reward devaluation experiment is not included in this pool; the same analysis for that experiment, together with representative single-subject data, is shown in Figure S5. ***a– c***, Press rate as a joint function of food-port occupancy decile and the duration of the analysis window, computed separately for the intertrial interval (a, ITI), pre-CS (b), and CS (c) epochs. Press rate declines as occupancy increases in every epoch, confirming that the two responses do compete when both are expressed. Color scales are set independently for each panel. ***d, e***, Press rate across port-occupancy deciles for a 10-s (d) and 1-s (e) analysis window, by epoch. Dotted gray, ITI; dashed green, pre-CS; solid purple, CS; horizontal dashed lines give each epoch’s overall mean. Bar histograms beneath each panel give the proportion of observations falling in each occupancy bin, pooled across epochs. The great majority of cue time is spent at low port occupancy: in the 10-s window, 83% of observations fall in the lowest occupancy decile, and the decline in press rate with increasing occupancy is confined to the sparsely populated upper deciles. Competition is thus a real but local phenomenon, expressed on a small minority of observations. Whether competition can constrain pressing also depends on how much of the available time rats actually used, which we quantified as utilization: the proportion of the cue period not spent in the food port during which a rat was engaged in a bout of lever pressing. Utilization averaged 5.1% across experiments. Rats therefore left the great majority of the time available to them unused, and competition for time cannot have acted as a mechanical limit on how much they pressed. Every effect of interest computed both with and without opportunity adjustment is reported in Tables 1–4. Data presented as mean ± SEM.

Port occupancy was nevertheless greater during CS_100_ than during CS_30_, so we asked whether the effects of interest survived after controlling for this cue-specific imbalance in available response time. We recomputed cue-evoked pressing using only the time during which each rat was free to press, a deliberately conservative treatment because it credits every second of port occupancy to competition. The findings generally held (Tables 1–4). Cue-specific modulation of press rate across control conditions remained reliable (t(74) = 2.34, p = 0.022), as did the effect of dmPFC inhibition (F(1,13) = 9.11, p = 0.0099). The omnibus devaluation effect was attenuated on the adjusted measure (F(1,22) = 3.17, p = 0.089; Table 2; see also Figure S5), but the planned comparison in the fully predicted (100%) group remained significant (t(11) = −2.40, p = 0.035).

Competition between these responses therefore does occur, but only on the minority of trials in which food-port occupancy is high, and it does not account for the cue-specific regulation of instrumental performance reported here. This complements other recent findings indicating that the PIT assay used here involves flexible regulation of the strategy used to pursue rewards and is not the byproduct of a simple response competition mechanism (Marshall and Ostlund, 2018; Marshall et al., 2020; Marshall et al., 2023; Malvaez et al., 2026).

## Discussion

The current study examined the role of the dmPFC in the flexible regulation of cue-motivated behavior when imminent reward is predicted. First, we show that rats rely on a goal-directed assessment of the predicted reward when deciding whether to withhold instrumental reward seeking, implying the use of a top-down cognitive control process. Second, we show that cues for imminent reward phasically stimulate Ca^2+^ activity in excitatory dmPFC neurons, and that this activity scales with the likelihood of upcoming reward and tracks whether reward seeking is expressed on a given trial. Finally, we show that the ability to use imminent reward predictions to flexibly regulate instrumental performance can be impaired by inhibiting the dmPFC, establishing a causal role for this region.

Reward-predictive cues shape the way rewards are pursued. Cues signaling that rewards are scarce, uncertain, or delayed promote exploratory, general search behaviors, including instrumental reward-seeking actions, whereas cues that signal imminent reward trigger a shift from general to focal search activities directed at the reward retrieval site (Konorski, 1967; Bindra, 1974; Timberlake, 1994). It has been argued that imminent reward cues trigger this shift through a top-down, cognitive control process (Ostlund and Marshall, 2021). This view assumes that cues for imminent reward acquire strong incentive motivational properties but do not invigorate instrumental performance because that behavior is actively suppressed in favor of the more advantageous goal-approach response. We therefore sought to expose this latent influence by removing the reason for its suppression: devaluing the predicted reward. Note first that cues signaling sparse reward availability retain their full capacity to stimulate instrumental performance even if the predicted reward has been devalued (Rescorla, 1994; Holland, 2004), indicating that this excitatory influence is mediated by a general motivational process rather than one that involves a cognitive assessment of expected rewards. By contrast, the current results suggest that both types of process are engaged by cues that signal imminent reward. Rats increased, rather than decreased, their instrumental lever-pressing when presented with a cue that signaled imminent delivery of a devalued reward. We also found that shifting rats from hunger to general satiety disrupted the use of expected reward probability to regulate instrumental performance. Both manipulations therefore implicate reward value in this regulation, but they differ in what they expose. Selectively devaluing the predicted reward revealed the cue’s latent motivational influence, whereas general satiety did not, and it also lowered baseline pressing. This likely reflects the known attenuating effect of general satiety on both the nonspecific motivational influence of reward-paired cues (general PIT) (Corbit et al., 2007) and instrumental response vigor, making this manipulation harder to attribute to a single process. Together these findings indicate that imminent reward cues trigger a real-time appraisal of the predicted reward, which we suggest is used to assess the value of withholding instrumental performance in favor of goal approach.

The dmPFC is essential for the flexible control of motivated behavior (Ragozzino, 2007; Bissonette and Roesch, 2017). Leading theories of dmPFC function assign it a central role in resolving conflict between competing action tendencies (Shenhav et al., 2013; Clairis and Lopez-Persem, 2023). Consistent with this, the dmPFC is preferentially activated by stimuli that elicit conflict, such as a signal to inhibit a prepotent response (MacDonald et al., 2000; Botvinick et al., 2001; Braver et al., 2001; Kerns et al., 2004). Theories of this kind separate two components of control: specifying how much control a situation warrants, and implementing it, with the dorsal anterior cingulate assigned the former role (Shenhav et al., 2013). A signal that specifies the current demand for control should scale with the strength of the impulse to be overcome, and should, therefore, be most pronounced on trials where that impulse is most likely to prevail. This is the case for the dorsal anterior cingulate, which is engaged both by high-conflict trials and by trials on which control fails (Braver et al., 2001). The activity reported here fits that profile and indicates that the dmPFC may play a similar role in resolving Pavlovian-instrumental conflict triggered by cues predicting imminent reward delivery. We found that dmPFC Ca^2+^ activity was phasically elevated by these cues, largely within the first few seconds after CS onset, and encoded the probability of upcoming reward, a critical factor determining the value of shifting from instrumental to goal-approach behavior. While similar results have been observed in Pavlovian (stimulus-reward) learning (Otis et al., 2017; Ottenheimer et al., 2023), the current study found evidence that this cue-elicited dmPFC Ca^2+^ activity varied with the cue’s influence on instrumental performance. The high-probability cue elicited a large increase in dmPFC Ca^2+^ activity that was heightened further on trials in which rats went on to press the lever. The low-probability cue elicited weaker dmPFC Ca^2+^ activity that did not differ between press and no-press trials. We take pressing during the high-probability cue to mark the trials on which the motivational impulse was strongest, and these were the trials on which the dmPFC was most engaged, as expected of a signal that specifies the momentary demand for control. The absence of this relationship during the low-probability cue, which poses no comparable control problem, argues against a general arousal or response coding account.

We also found that this ability to regulate instrumental performance using reward predictions depends on the dmPFC. Inhibiting dmPFC neurons abolished the cue-specific regulation of lever pressing. Rats pressed at comparable rates during cues signaling a high and a low probability of imminent reward. This disruptive effect of dmPFC inhibition was specific to instrumental performance, sparing rats’ ability to use cue-elicited reward predictions to differentially control their conditioned goal-approach behavior. CNO did reduce overall food-port occupancy in these rats, but it left the cue-specific structure of that behavior intact, so the deficit cannot be attributed to impairments in cue detection, discrimination, memory retrieval, or gross motor control. This specificity, together with the finding that rats had ample opportunity to press during both cues, also argues against the possibility that cue-specific differences in pressing reflect competition for time with food-port approach.

Although the present experiments did not target specific dmPFC output pathways, the dmPFC projection to the nucleus accumbens is well-situated to contribute to this form of regulation. The nucleus accumbens is known to mediate the excitatory effects of reward-paired cues on instrumental performance (Corbit and Balleine, 2015), with local dopamine transmission in this structure playing a key role (Wyvell and Berridge, 2000; Lex and Hauber, 2008; Wassum et al., 2013; Halbout et al., 2019). Although nucleus accumbens dopamine transmission is required for the excitatory effects of reward-paired cues in conventional PIT studies (Lex and Hauber, 2008; Halbout et al., 2019), recent work using the same probabilistic PIT task variant applied here to study Pavlovian-instrumental conflict found that cues for imminent reward elicit phasic nucleus accumbens dopamine release that encodes reward probability (Malvaez et al., 2026). However, unlike the dmPFC Ca^2+^ signal reported here, cue-evoked nucleus accumbens dopamine responses were inversely correlated with instrumental performance and positively correlated with goal approach. Moreover, optogenetic inhibition of nucleus accumbens dopamine terminals during that cue amplified instrumental performance while attenuating goal approach. We therefore suggest that dmPFC projections to the nucleus accumbens or its mesolimbic dopamine afferents may modulate dopamine responses to imminent reward cues to bias control away from instrumental seeking and toward goal approach, although we have not tested this directly. The dmPFC projection to the paraventricular nucleus of the thalamus is an equally plausible route, having already been implicated in the top-down control of cue-directed approach (Campus et al., 2019). Whether either pathway contributes to the regulation of instrumental reward seeking when imminent reward is predicted has yet to be examined. Cue-evoked dmPFC activity encoded reward probability and showed prediction error-like features at the expected time of delivery, but whether it also encodes the incentive value of the predicted reward, the dimension shown here to control instrumental regulation, is not yet known. Another limitation of our study is that only the satiety experiment included both sexes, leaving open questions about whether our other findings generalize across sexes.

The present findings contribute to a growing literature implicating Pavlovian-instrumental conflict as a fundamental challenge for behavioral regulation that recruits prefrontal-striatal control mechanisms across species, including human work on how predictive cues generate prepotent response biases that must be overcome through prefrontal-dependent control (Dorfman and Gershman, 2019; Gershman et al., 2020). Studies using this paradigm have also shown that the ability to adaptively resolve Pavlovian-instrumental conflict is altered in children (Raab and Hartley, 2020) and obsessive compulsive disorder (Peng et al., 2022), and is associated with problematic alcohol consumption (Chen et al., 2020; Chen et al., 2023). Similarly, preclinical studies have shown that both adolescent (Marshall et al., 2020) and psychostimulant-sensitized rats (Marshall and Ostlund, 2018) display abnormally vigorous instrumental performance in response to imminent reward cues, a profile resembling the loss of cue-specific regulation produced by dmPFC inhibition here. Dysfunction of this prefrontal control mechanism may therefore represent a shared point of vulnerability in the development of impulsive, risky reward seeking.

## Table Titles

Table 1. Cue-specific regulation of instrumental pressing is intact under unperturbed conditions (pooled control-only sample).

Table 2. Reward devaluation releases cue-evoked pressing.

Table 3. Satiety disrupts cue-specific regulation of instrumental pressing (State × Cue interaction).

Table 4. dmPFC inhibition disrupts cue-specific regulation of instrumental pressing (Drug × Cue interaction).

Opportunity adjustment expresses press rate per second of time not spent occupying the food port, so that cue-evoked differences in port occupancy cannot mechanically produce a difference in press rate. Unadjusted and adjusted analyses use different dependent variables and are both reported; they are not competing estimates of the same quantity. Adjusted press rate is computed separately for each subject and cue as a ratio of sums, dividing total presses by total time not spent in the food port across that cue’s trials, and is applied to the CS and pre-CS periods alike (Figure 5).

## Author contributions

[B.H., K.M.W. and S.B.O] designed research; [B.H., C.H., N.R., G.A., N.N. and S.B.O.] performed research; [B.H. and S.B.O.] analyzed data; [B.H. and S.B.O] wrote the first draft of the paper; [B.H., K.M.W. and S.B.O.] edited the paper.

## Acknowledgments

This work was supported by NIMH grant MH126285 (SBO and KMW) and NIDA grant DA064341 (SBO). Claude (Anthropic) was used to assist with editing this manuscript for clarity, grammar, and journal formatting.

## References

Azrin N, Hake D (1969) Positive conditioned suppression: conditioned suppression using positive reinforcers as the unconditioned stimuli 1. J Exp Anal Behav 12:167–173.

Bindra D (1974) A motivational view of learning, performance, and behavior modification. Psychological Review 81:199–213.

Bissonette GB, Roesch MR (2017) Neurophysiology of rule switching in the corticostriatal circuit. Neuroscience 345:64–76.

Botvinick MM, Braver TS, Barch DM, Carter CS, Cohen JD (2001) Conflict monitoring and cognitive control. Psychol Rev 108:624–652.

Braver TS, Barch DM, Gray JR, Molfese DL, Snyder A (2001) Anterior cingulate cortex and response conflict: effects of frequency, inhibition and errors. Cereb Cortex 11:825–836.

Campus P, Covelo IR, Kim Y, Parsegian A, Kuhn BN, Lopez SA, Neumaier JF, Ferguson SM, Solberg Woods LC, Sarter M (2019) The paraventricular thalamus is a critical mediator of top-down control of cue-motivated behavior in rats. Elife 8:e49041.

Cardinal RN, Parkinson JA, Marbini HD, Toner AJ, Bussey TJ, Robbins TW, Everitt BJ (2003) Role of the anterior cingulate cortex in the control over behavior by Pavlovian conditioned stimuli in rats. Behav Neurosci 117:566.

Chen H, Belanger MJ, Garbusow M, Kuitunen-Paul S, Huys QJM, Heinz A, Rapp MA, Smolka MN (2023) Susceptibility to interference between Pavlovian and instrumental control predisposes risky alcohol use developmental trajectory from ages 18 to 24. Addiction Biology 28:e13263.

Chen H, Nebe S, Mojtahedzadeh N, Kuitunen-Paul S, Garbusow M, Schad D, Rapp M, Huys Q, Heinz A, Smolka M (2020) Susceptibility to interference between Pavlovian and instrumental control is associated with early hazardous alcohol use. Addiction Biology 26.

Clairis N, Lopez-Persem A (2023) Debates on the dorsomedial prefrontal/dorsal anterior cingulate cortex: insights for future research. Brain 146:4826–4844.

Corbit LH, Balleine BW (2003) The role of prelimbic cortex in instrumental conditioning. Behavioural brain research 146:145–157.

Corbit LH, Balleine BW (2015) Learning and motivational processes contributing to Pavlovian– instrumental transfer and their neural bases: dopamine and beyond. Curr Top Behav Neuro:259–289.

Corbit LH, Janak PH, Balleine BW (2007) General and outcome-specific forms of Pavlovian-instrumental transfer: the effect of shifts in motivational state and inactivation of the ventral tegmental area. Eur J Neurosci 26:3141–3149.

Crombag HS, Galarce EM, Holland PC (2008) Pavlovian influences on goal-directed behavior in mice: The role of cue-reinforcer relations. Learn Memory 15:299–303.

Dorfman H, Gershman S (2019) Controllability governs the balance between Pavlovian and instrumental action selection. Nature Communications 10.

Estes WK (1943) Discriminative conditioning. I. A discriminative property of conditioned anticipation. Journal of Experimental Psychology 32:150.

Gershman S, Guitart-Masip M, Cavanagh J (2020) Neural signatures of arbitration between Pavlovian and instrumental action selection. PLoS Computational Biology 17.

Halbout B, Hutson C, Wassum KM, Ostlund SB (2022) Dorsomedial prefrontal cortex activation disrupts Pavlovian incentive motivation. Frontiers in Behavioral Neuroscience Volume 16–2022.

Halbout B, Marshall AT, Azimi A, Liljeholm M, Mahler SV, Wassum KM, Ostlund SB (2019) Mesolimbic dopamine projections mediate cue-motivated reward seeking but not reward retrieval in rats. Elife 8.

Holland PC (2004) Relations between Pavlovian-instrumental transfer and reinforcer devaluation. J Exp Psychol Anim Behav Process 30:104–117.

Homayoun H, Moghaddam B (2009) Differential representation of Pavlovian-instrumental transfer by prefrontal cortex subregions and striatum. Eur J Neurosci 29:1461–1476.

Howland JG, Ito R, Lapish CC, Villaruel FR (2022) The rodent medial prefrontal cortex and associated circuits in orchestrating adaptive behavior under variable demands. Neuroscience & Biobehavioral Reviews 135:104569.

Jean-Richard-dit-Bressel P, Clifford CWG, McNally GP (2020) Analyzing Event-Related Transients: Confidence Intervals, Permutation Tests, and Consecutive Thresholds. Frontiers in Molecular Neuroscience Volume 13–2020.

Keevers LJ, Jean-Richard-Dit-Bressel P (2025) Obtaining artifact-corrected signals in fiber photometry via isosbestic signals, robust regression, and dF/F calculations. Neurophotonics 12:025003.

Kerns JG, Cohen JD, MacDonald AW, 3rd, Cho RY, Stenger VA, Carter CS (2004) Anterior cingulate conflict monitoring and adjustments in control. Science 303:1023–1026.

Konorski J (1967) Integrative activity of the brain.

Lex A, Hauber W (2008) Dopamine D1 and D2 receptors in the nucleus accumbens core and shell mediate Pavlovian-instrumental transfer. Learn Memory 15:483–491.

MacDonald AW, 3rd, Cohen JD, Stenger VA, Carter CS (2000) Dissociating the role of the dorsolateral prefrontal and anterior cingulate cortex in cognitive control. Science 288:1835–1838.

Malvaez M, Suarez A, Griffin NK, Ramírez-Armenta K, Ostlund SB, Wassum KM (2026) Dopamine Supports Reward Prediction to Shape Reward-Pursuit Strategy. The Journal of Neuroscience 46:e1636252026.

Marshall AT, Ostlund SB (2018) Repeated cocaine exposure dysregulates cognitive control over cue-evoked reward-seeking behavior during Pavlovian-to-instrumental transfer. Learn Memory 25:399–409.

Marshall AT, Munson CN, Maidment NT, Ostlund SB (2020) Reward-predictive cues elicit excessive reward seeking in adolescent rats. Dev Cogn Neurosci 45:100838.

Marshall AT, Halbout B, Munson CN, Hutson C, Ostlund SB (2023) Flexible control of Pavlovian-instrumental transfer based on expected reward value. J Exp Psychol Anim Learn Cogn 49:14–30.

McLaughlin AE, Diehl GW, Redish AD (2021) Potential roles of the rodent medial prefrontal cortex in conflict resolution between multiple decision-making systems. International review of neurobiology 158:249–281.

Miczek KA, Grossman SP (1971) Positive conditioned suppression: Effects of CS duration 1. J Exp Anal Behav 15:243–247.

Ostlund SB, Marshall AT (2021) Probing the role of reward expectancy in Pavlovian-instrumental transfer. Current Opinion in Behavioral Sciences 41:106–113.

Otis JM, Namboodiri VM, Matan AM, Voets ES, Mohorn EP, Kosyk O, McHenry JA, Robinson JE, Resendez SL, Rossi MA, Stuber GD (2017) Prefrontal cortex output circuits guide reward seeking through divergent cue encoding. Nature 543:103–107.

Ottenheimer DJ, Hjort MM, Bowen AJ, Steinmetz NA, Stuber GD (2023) A stable, distributed code for cue value in mouse cortex during reward learning. eLife 12:RP84604.

Peng Z, He L, Wen R, Verguts T, Seger CA, Chen Q (2022) Obsessive-compulsive disorder is characterized by decreased Pavlovian influence on instrumental behavior. PLoS Computational Biology 18:e1009945.

Raab HA, Hartley CA (2020) Adolescents exhibit reduced Pavlovian biases on instrumental learning. Sci Rep 10:15770.

Ragozzino ME (2007) The contribution of the medial prefrontal cortex, orbitofrontal cortex, and dorsomedial striatum to behavioral flexibility. Ann N Y Acad Sci 1121:355–375.

Rescorla RA (1994) Transfer of instrumental control mediated by a devalued outcome. Animal Learning & Behavior 22:27–33.

Robinson T, Carr C, Kawa A (2018) The propensity to attribute incentive salience to drug cues and poor cognitive control combine to render sign-trackers susceptible to addiction. Sign-tracking and drug addiction (Vol A) 10.

Rudebeck PH, Izquierdo A (2022) Foraging with the frontal cortex: A cross-species evaluation of reward-guided behavior. Neuropsychopharmacology 47:134–146.

Sharpe MJ, Stalnaker T, Schuck NW, Killcross S, Schoenbaum G, Niv Y (2019) An integrated model of action selection: distinct modes of cortical control of striatal decision making. Annual review of psychology 70:53–76.

Shenhav A, Botvinick Matthew M, Cohen Jonathan D (2013) The Expected Value of Control: An Integrative Theory of Anterior Cingulate Cortex Function. Neuron 79:217–240.

Tan SYS, Shen MH, Keevers LJ, Williams-Spooner M, McNally GP, Killcross S, Jean-Richard-dit-Bressel P (2026) Disinhibition of ventral tegmental area during initial punishment learning causes enduring punishment insensitivity. Neuropsychopharmacology.

Timberlake W (1988) Evolution, behavior systems, and “self-control”: The fit between organism and test environment. Behavioral and Brain Sciences 11:694–695.

Timberlake W (1994) Behavior systems, associationism, and Pavlovian conditioning. Psychonomic Bulletin & Review 1:405–420.

Wassum KM, Ostlund SB, Loewinger GC, Maidment NT (2013) Phasic Mesolimbic Dopamine Release Tracks Reward Seeking During Expression of Pavlovian-to-Instrumental Transfer. Biological Psychiatry 73:747–755.

Wyvell CL, Berridge KC (2000) Intra-accumbens amphetamine increases the conditioned incentive salience of sucrose reward: enhancement of reward “wanting” without enhanced “liking” or response reinforcement. Journal of Neuroscience 20:8122–8130.

