## Supplementary figures and images for "The Dorsomedial Prefrontal Cortex Uses Reward Predictions to Regulate How Rewards Are Pursued"

### Supplemental Figure 1

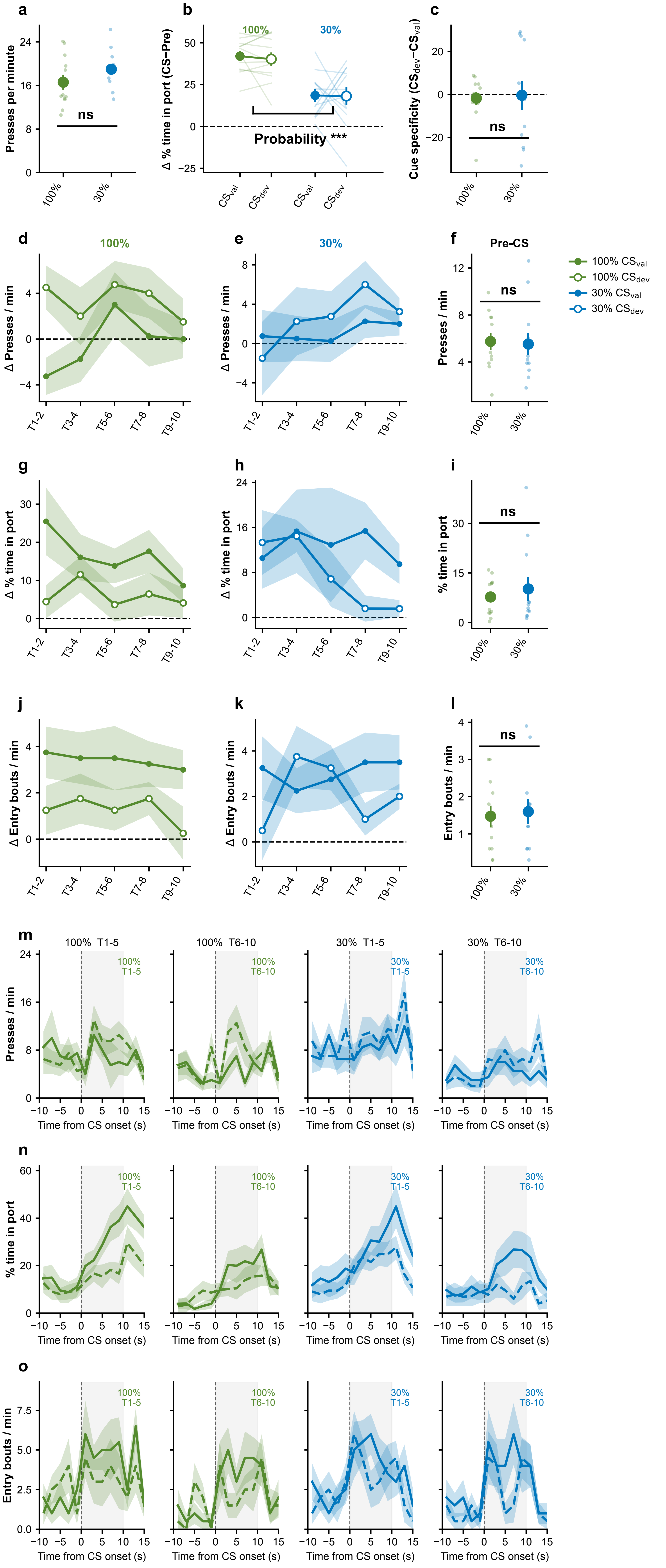

### Supplemental Figure 2

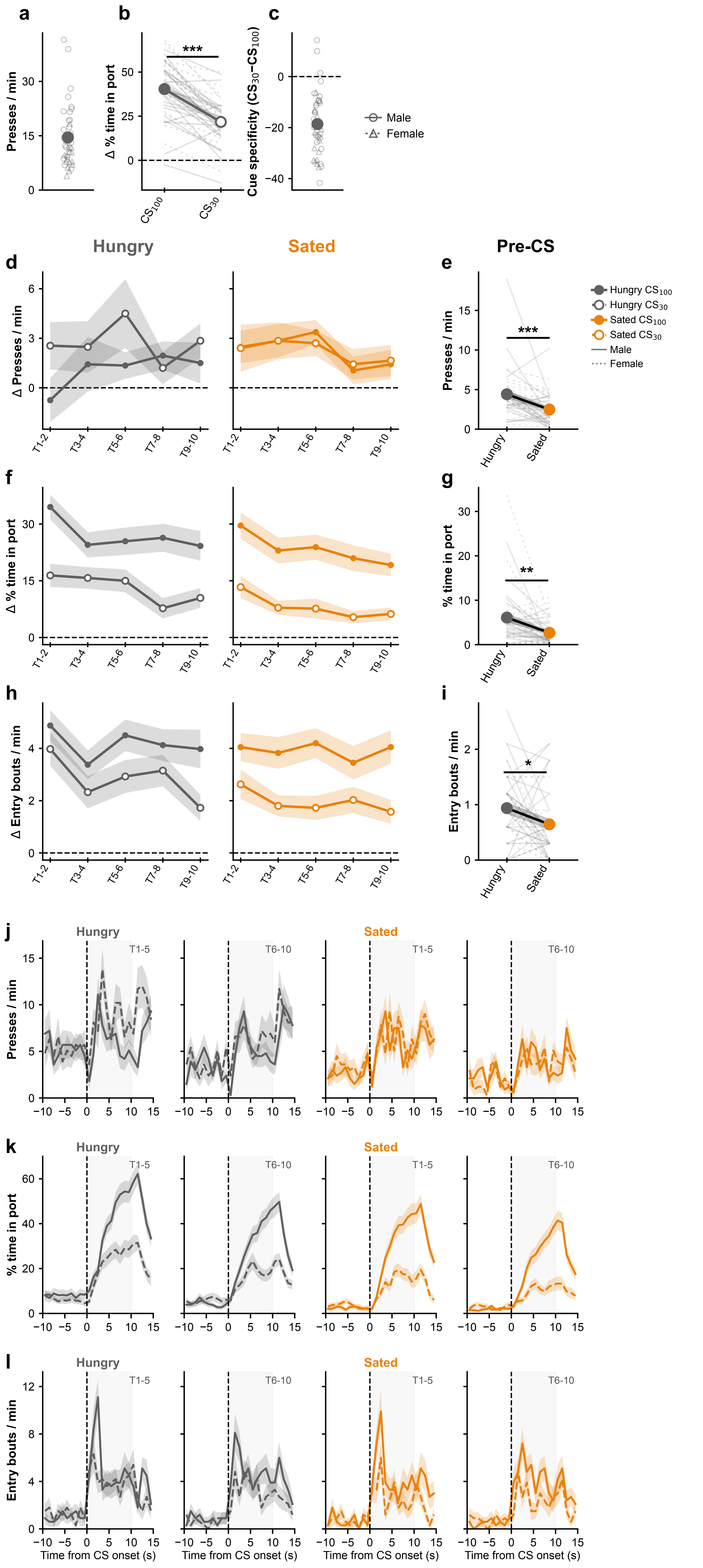

### Supplemental Figure 3

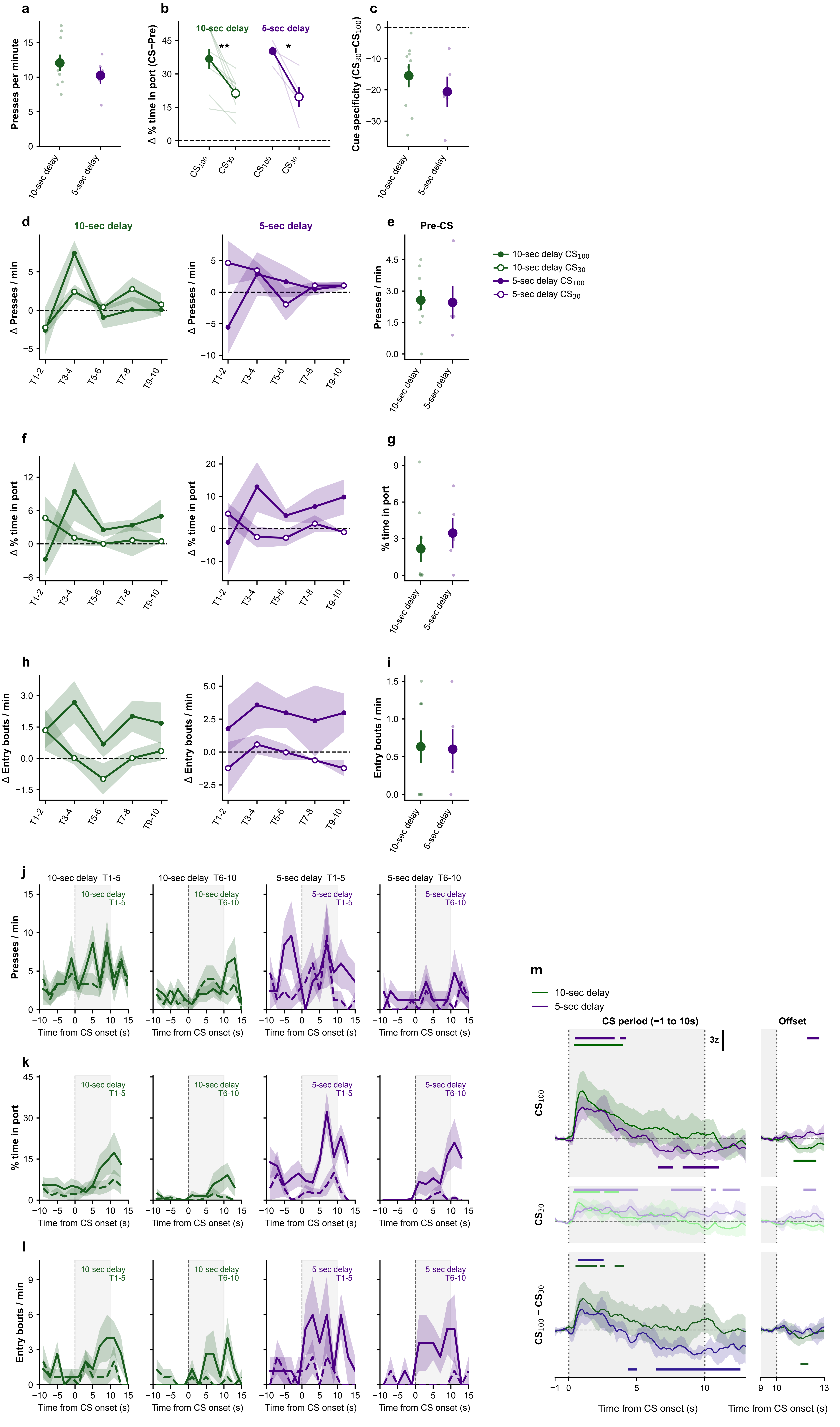

### Supplemental Figure 4

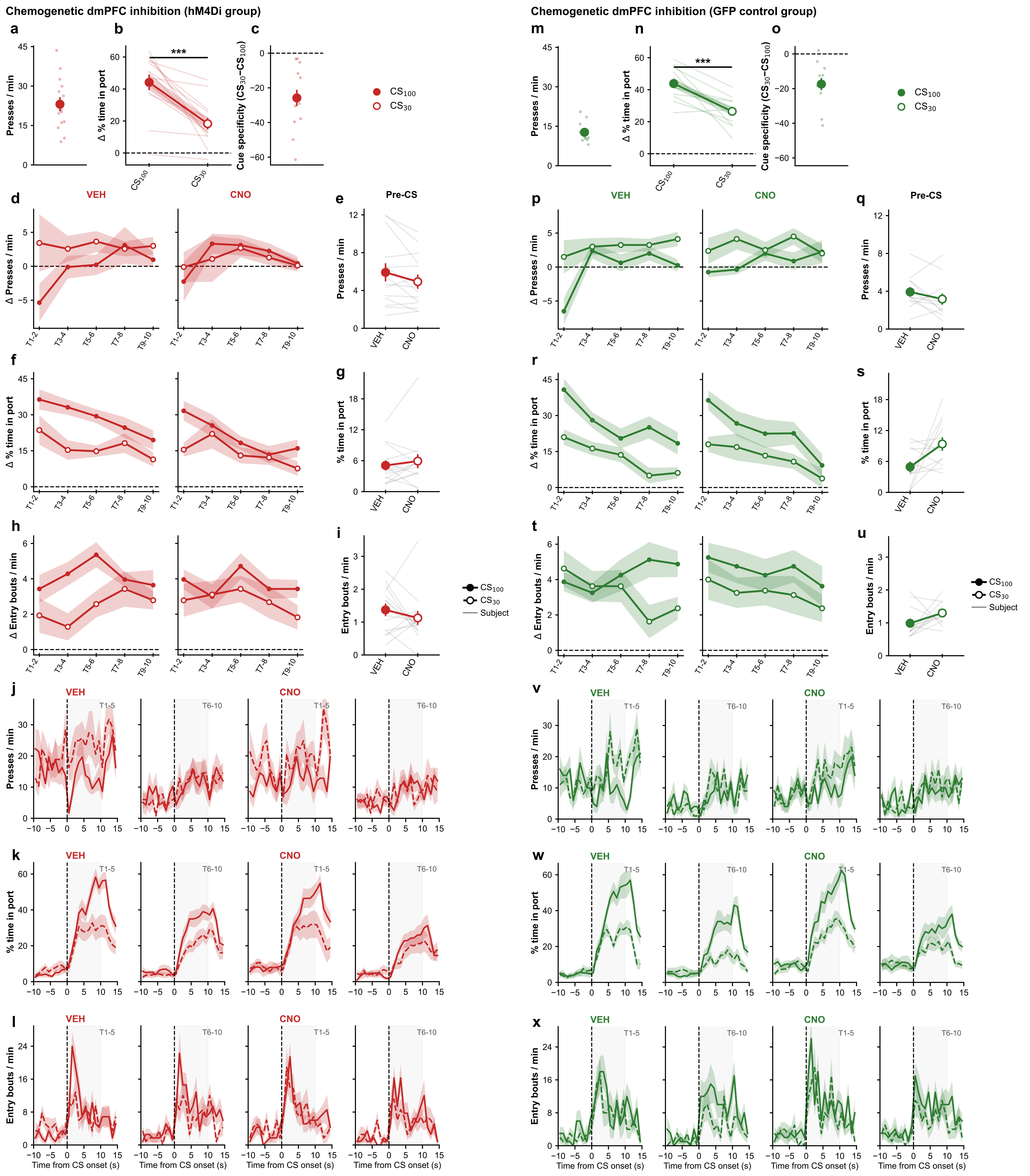

### Supplemental Figure 5

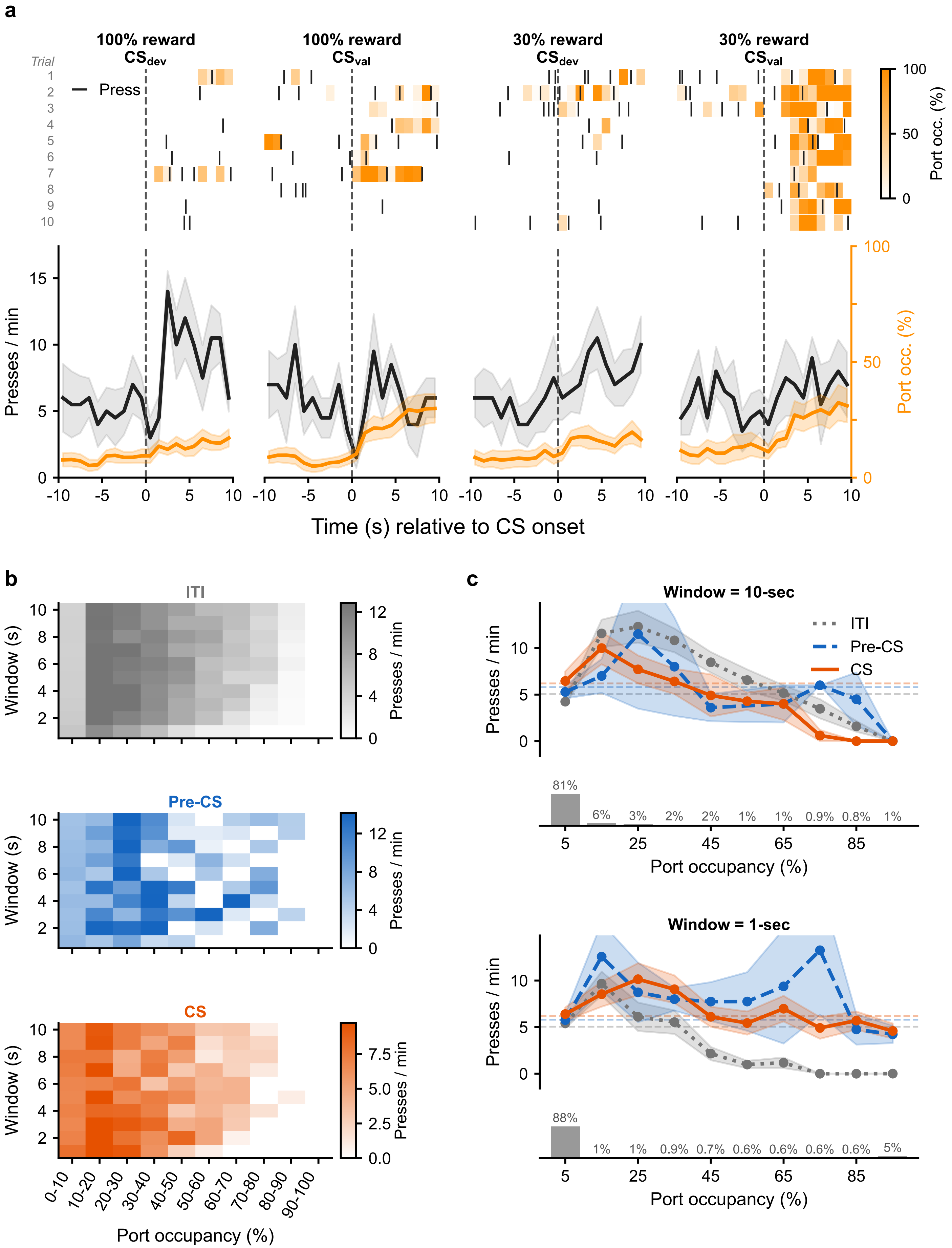
