## Supplementary material for "The Dorsomedial Prefrontal Cortex Uses Reward Predictions to Regulate How Rewards Are Pursued": Tables

Main Tables

Table 1. Cue-specific regulation of instrumental pressing is intact under unperturbed conditions (pooled control-only sample).

| n | Opportunity-adjustment | $CS_{100}$ ΔPress rate (± SEM) | $CS_{30}$ ΔPress rate (± SEM) | Effect ± SE | Test | p | Effect size [95% CI] |
|---|---|---|---|---|---|---|---|
| 75 | Unadjusted | 0.28 ± 0.57 | 2.83 ± 0.76 | 2.54 ± 0.64 | t(74) = 3.962 | 0.0002 | *d* = 0.457 [0.218, 0.694] |
| 75 | Adjusted | 2.65 ± 0.72 | 4.41 ± 0.80 | 1.76 ± 0.75 | t(74) = 2.339 | 0.0220 | *d* = 0.270 [0.039, 0.500] |

Table 2. Reward devaluation releases cue-evoked pressing.

| n | Term | Opportunity-adjustment | ΔPress rate Val ± SEM | ΔPress rate Dev ± SEM | Effect ± SE | Test | p | Effect size [95% CI] |
|---|---|---|---|---|---|---|---|---|
| 24 | Omnibus cue (devaluation effect, all groups) | Unadjusted | | | −2.55 ± 1.09 | F(1,22) = 5.537 | 0.0280 | $\eta^2_p$ = 0.201 [0.000, 0.447] |
| 24 | Omnibus cue (devaluation effect, all groups) | Adjusted | | | −2.32 ± 1.29 | F(1,22) = 3.168 | 0.0889 | $\eta^2_p$ = 0.126 [0.000, 0.374] |
| 12 | Val vs Dev @ 100% group | Unadjusted | −0.35 ± 0.80 | 3.35 ± 1.04 | −3.70 ± 1.08 | t(11) = −3.426 | 0.0057 | $d_z$ = −0.989 [−1.672, −0.277] |
| 12 | Val vs Dev @ 100% group | Adjusted | 0.79 ± 0.96 | 4.12 ± 1.09 | −3.33 ± 1.39 | t(11) = −2.401 | 0.0352 | $d_z$ = −0.693 [−1.315, −0.047] |
| 12 | Val vs Dev @ 30% group | Unadjusted | 1.15 ± 0.91 | 2.55 ± 1.36 | −1.40 ± 1.88 | t(11) = −0.745 | 0.4719 | $d_z$ = −0.215 [−0.783, 0.362] |
| 12 | Val vs Dev @ 30% group | Adjusted | 2.43 ± 1.37 | 3.75 ± 1.43 | −1.31 ± 2.21 | t(11) = −0.594 | 0.5644 | $d_z$ = −0.172 [−0.738, 0.403] |

Table 3. Satiety disrupts cue-specific regulation of instrumental pressing (State × Cue interaction).

| n | Opportunity-adjustment | Test | p | Effect size [95% CI] |
|---|---|---|---|---|
| 40 | Unadjusted | F(1,39) = 5.442 | 0.0249 | $\eta^2_p$ = 0.122 [0.000, 0.315] |
| 40 | Adjusted | F(1,39) = 4.134 | 0.0489 | $\eta^2_p$ = 0.096 [0.000, 0.284] |

Table 4. dmPFC inhibition disrupts cue-specific regulation of instrumental pressing (Drug × Cue interaction).

| n | Group | Opportunity-adjustment | Test | p | Effect size [95% CI] |
|---|---|---|---|---|---|
| 14 | hM4Di | Unadjusted | $F(1,13) = 15.113$ | 0.0019 | $\eta^2_p = 0.538$ [0.113, 0.725] |
| 14 | hM4Di | Adjusted | $F(1,13) = 9.113$ | 0.0099 | $\eta^2_p = 0.412$ [0.031, 0.647] |
| 12 | GFP | Unadjusted | $F(1,11) = 1.040$ | 0.3297 | $\eta^2_p = 0.086$ [0.000, 0.409] |
| 12 | GFP | Adjusted | $F(1,11) = 0.670$ | 0.4306 | $\eta^2_p = 0.057$ [0.000, 0.374] |

Opportunity adjustment expresses press rate per second of time not spent occupying the food port, so that cue-evoked differences in port occupancy cannot mechanically produce a difference in press rate. Unadjusted and adjusted analyses use different dependent variables and are both reported; they are not competing estimates of the same quantity. Adjusted press rate is computed separately for each subject and cue as a ratio of sums, dividing total presses by total time not spent in the food port across that cue's trials, and is applied to the CS and pre-CS periods alike (Figure 5).
